# Secretome analysis of human and rat pancreatic islets co-cultured with adipose-derived stromal cells reveals a signature with enhanced regenerative capacities

**DOI:** 10.1101/2024.11.01.621481

**Authors:** Erika Pinheiro-Machado, Bart J. de Haan, Marten A. Engelse, Alexandra M. Smink

## Abstract

Pancreatic islet transplantation (PIT), a promising treatment for Type 1 Diabetes (T1D), encounters challenges in the pre- and post-transplantation phases. Co-culturing or co-transplantation of islets with mesenchymal stromal cells (MSC), known for their regenerative properties, emerged as a potential solution and was shown to increase islet function and improve PIT outcomes. This study explored the changes in the islets’ secretion signature (secretome) when co-cultured with adipose-derived stromal cells (ASC), an MSC subtype. The secretome profile of islets and co-cultures under various stressors, i.e., cytokines, high glucose, hypoxia, and a combination of hypoxia and high glucose, was investigated. The results shed light on the potential mechanisms through which ASC support islets’ functional survival. Co-culturing pancreatic islets with ASC induced substantial proteomic changes, impacting pathways crucial for energy metabolism, angiogenesis, extracellular matrix organization, and immune responses. The analysis of key signaling molecules (VEGF, PDGF, bFGF, Collagen I alpha 1, IL-1α, and IL-10) revealed alterations influenced by the culturing conditions and the presence of the ASC. *In vitro* functional assays using the secretomes also demonstrated their potential to differentially influence angiogenic processes, enhance collagen deposition, and modulate the immune system based on the conditions in which they were generated. These findings offer valuable insights into the potential of ASC co-culturing to address challenges in PIT, paving the way for enhanced therapeutic interventions in T1D and regenerative medicine.

**Highlights:**

- Both *in-silico* and *in-vitro* data support that co-culturing pancreatic islets with ASC enhances islet function.
- Co-culturing islets with ASC induces changes in the secretome, impacting pathways related to energy metabolism, angiogenesis, extracellular matrix organization, and immune response.
- Key signaling molecules, including VEGF, PDGF, bFGF, Collagen I alpha 1, IL-1α, and IL-10, are differentially affected by various co-culturing conditions.

## 1. Introduction

Pancreatic islet transplantation (PIT), a treatment option for Type 1 Diabetes (T1D), faces obstacles in both pre-[1, 2] and post-transplantation phases [3, 4]. Its long-term success is hindered by challenges such as insufficient vascularization, extracellular matrix (ECM) damage, weak engraftment, and immunological responses [3–5]. Addressing these issues is crucial for enhancing the functional survival of transplanted islets and extending the benefits of this treatment to a larger number of patients.

To overcome the loss of islet quality and poor survival rates, researchers have explored co-culturing or co-transplanting islets with mesenchymal stromal cells (MSC) [5–7]. MSC are a subset of heterogeneous non-hematopoietic fibroblast-like cells with multi-lineage differentiation capacity [8, 9]. These multipotent MSC can be obtained from various sources, such as the adipose tissue (adipose-derived stromal cells (ASC)), contributing to tissue repair through migration and secretion of biologically active molecules with anti-inflammatory, immunomodulatory, and angiogenic properties [10].

Several studies have demonstrated the effectiveness of co-culturing/co-transplantation islets with MSC, including ASC, showing enhanced insulin secretion, reduced beta cell apoptosis, and increased beta cell mass [11–13]. The positive results were shown both *in vitro* and in T1D animal models [14–17]. However, the mechanism behind the positive PIT effects of MSC co-culturing or transplantation with islets remains unknown. This mechanism might be either by direct contact between islets and MSC or by producing various biologically active molecules (the secretome). This study investigates the composition of the secretome and their angiogenic, ECM modulating, and immunomodulatory capacities as these will improve the microenvironment and islet quality and survival rates following PIT. By examining changes in the factors secreted by islets alone vs. islets co-cultured with ASC, we aim to uncover key components and pathways influencing islet function and survival following co-culture with ASC. This investigation into the secretome is pivotal for a comprehensive understanding of the dynamics driving the therapeutic benefits of ASC co-cultured with islets, offering insights for future refinement of therapeutic strategies.

## 2. Methods

### 2.1. Experimental design

In this study, the changes in the secretion profile of human and rat pancreatic islets cultured alone or in co-culture with human or rat adipose-derived stromal cells (h-prASC or r-prASC) were explored when subjected to various conditions designed to induce a different type of stress, mimicking challenges encountered during isolation and PIT. These conditions included normoxia – as the baseline control –, exposure to cytokines, high glucose, hypoxia, and a combination of hypoxia and high glucose (Fig. 1). Co-culturing was investigated in two distinct ratios of islets to ASC, 1:300 or 1:1000. Islets and co-cultures were subjected to these conditions for 72 hours (h). Ultimately, the supernatants (secretomes) were collected and analyzed using mass spectrometry, Luminex, and ELISA. The secretion profiles were characterized, and pathway enrichment was described. To validate the *in-silico* findings, we conducted *in vitro* functional investigations, focusing on the secretome’s capacity to stimulate critical processes such as angiogenesis, ECM deposition, and the attenuation of alloimmunity. These were achieved by performing a tube formation assay (TFA), Picrosirius Red staining, and a two-way mixed lymphocyte reaction (MLR) followed by an antibody-mediated cell-dependent cytotoxicity assay (CDC).

**Fig. 1.**
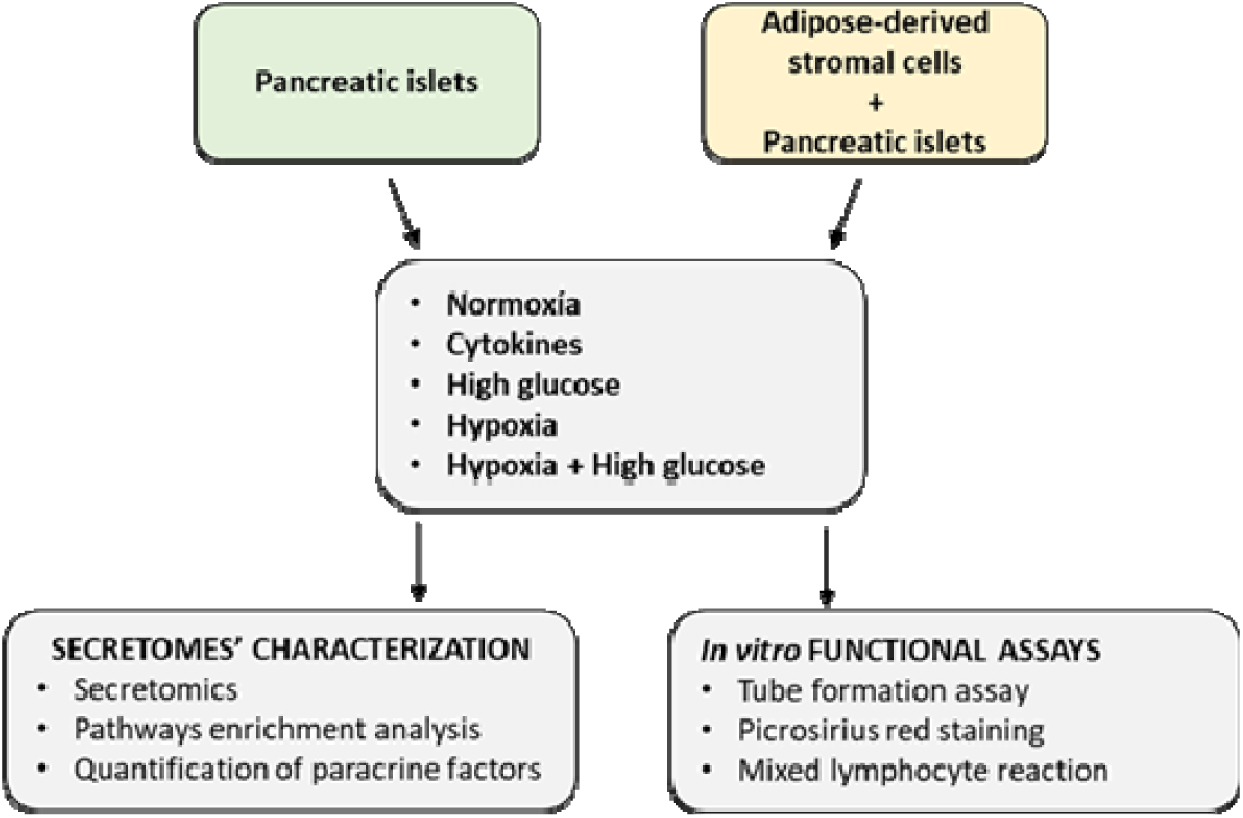
A summary of the experimental design.

### 2.2. Human and rat perirenal adipose tissue

This study used human and rat donors to obtain ASC, as previously described by Pinheiro-Machado, E. *et al.* (Pinheiro-Machado, E., et al. (2024) Culturing conditions dictate the composition and pathways enrichment of human and rat perirenal adipose-derived stromal cells’ secretomes. Stem Cell Reviews and Reports. Advanced online publication.). Human perirenal adipose tissue was procured from living kidney donors (n = 5) at the University Medical Center Groningen (UMCG; Groningen, The Netherlands). These samples were anonymously donated with informed consent after approval from the UMCG ethical review board. Maintained at 4°C, the human samples were processed within a 48-h time window. In turn, rat perirenal adipose tissue was collected from 7-9-week-old male Sprague-Dawley rats (n = 4; Envigo, Horst, The Netherlands). Ethical approval for tissue collection and procedures was obtained from the Dutch Central Committee on Animal Testing and the University of Groningen’s Animal Welfare Authority (AVD10500202115138).

### 2.3. Medium preparation for culturing and secretome collection

For the prASC isolation, a standard medium without serum (STD-M (−)) was used. STD-M (−) consisted of Dulbecco’s Modified Eagles Medium 4.5 g/L D-glucose (DMEM; Lonza, Walkersville, MD, USA) supplemented with 50 U/mL penicillin (Thermo Fisher Scientific, Bleiswijk, The Netherlands), 50 µg/mL streptomycin (Thermo Fisher Scientific), and 2 mM L-glutamine (Thermo Fisher Scientific). For the prASC and fibroblast culturing, a standard medium containing serum (STD-M (+)) was used. STD-M (+) contains DMEM 4.5 g/L D-glucose supplemented with 50 U/mL penicillin, 50 µg/mL streptomycin, and 2 mM L-glutamine, and 10% heat-inactivated Fetal Bovine Serum (FBS; Thermo Fisher Scientific). For culturing the islets, Connaught Medical Research Laboratories (CMRL) medium containing serum (CMRL-M (+)) was used. CMRL-M (+) consisted of CMRL medium (Thermo Fisher Scientific) supplemented with 8.3 mM glucose, 10% FBS, 2 mM GlutaMax supplement (Thermo Fisher Scientific), 50 U/mL penicillin, and 50 µg/mL streptomycin. For harvesting the normoxia and hypoxia-derived secretomes from the islets and co-cultures, CMRL without serum CMRL-M (−) was used. CMRL-M (−) consisted of CMRL medium supplemented with 8.3 mM glucose, 2 mM GlutaMax supplement, 50 U/mL penicillin, and 50 µg/mL streptomycin. The cytokines-derived secretomes were collected in CMRL-M (−) supplemented with 50 ng/mL interferon-gamma (IFN-γ) (ImmunoTools, Friesoythe, Germany), 21.5 ng/mL tumour necrosis factor-alpha (TNF-α) (ImmunoTools), and 0.25 ng/mL interleukin 1 beta (IL-1β) (ImmunoTools). Human and rat cytokines were obtained from ImmunoTools, and the same concentrations were used for both species. The high glucose and hypoxia + high glucose-derived secretomes were collected in CMRL-M (−) adjusted to 16.7 mM glucose. RPMI 1640 (Thermo Fisher Scientific) containing 4.5 g/L D-glucose and FBS (RPMI (+)) was used to culture human PBMC and rat splenocytes. RPMI (+) was supplemented with 50 U/mL penicillin, 50 µg/mL streptomycin, 2 mM L-glutamine, and 10% FBS. For the MLR, RPMI 1640 without serum (RPMI (−)) was used as a control.

### 2.4. prASC isolation and culture

Both rat and human prASC were isolated following the protocol previously described (Pinheiro-Machado, E., et al. (2024) Culturing conditions dictate the composition and pathways enrichment of human and rat perirenal adipose-derived stromal cells’ secretomes. Stem Cell Reviews and Reports. Advanced online publication.). Briefly, the tissue was washed, minced, and digested in a standard medium depleted of serum (STD-M (−)) containing NB 4 collagenase (0.5 mg/mL; Nordmark Biochemical, Uetersen, Germany) for 30 minutes (min) at 37°C. Cells were then separated from debris via centrifugation (500 x g, 5 min), and the resulting pellet was resuspended in STD-M (+). Resuspended cells were plated in T25 flasks for initial cell culture [Passage 0] at 37 °C and 5% CO_2_. After 24 h, debris and non-adherent cells were removed, and fresh STD-M (+) was added to the adherent cells. This medium was replaced every three days. At 80% confluence, cells were passaged by trypsinization. They were then expanded until passage 3 (h-prASC) or 2 (r-prASC) and stored in liquid nitrogen until they were needed for the co-culture experiments with rat and human islets. Both human and rat prASC have been fully characterized (Pinheiro-Machado, E., et al. (2024) Culturing conditions dictate the composition and pathways enrichment of human and rat perirenal adipose-derived stromal cells’ secretomes. Stem Cell Reviews and Reports. Advanced online publication.).

### 2.5. Human pancreatic islets

Human islets were procured either from Leiden University Medical Center (LUMC; Leiden, The Netherlands) or the European Consortium for Islet Transplantation (ECIT; Milan, Italy) [18, 19]. Detailed information about the donors and islets’ characteristics can be found in Table 1. These islets were assigned for research purposes as they did not meet the quality and/or quantity criteria necessary for clinical use. Research consents were obtained following the respective centers’ national regulations. Upon arrival at the UMCG, the islets were hand-picked and cultured in CMRL-M (+) for 24 h in an incubator set at 37 °C and 5% CO_2_. After acclimatization, these islets were cultured for secretome collection.

**Table 1.**
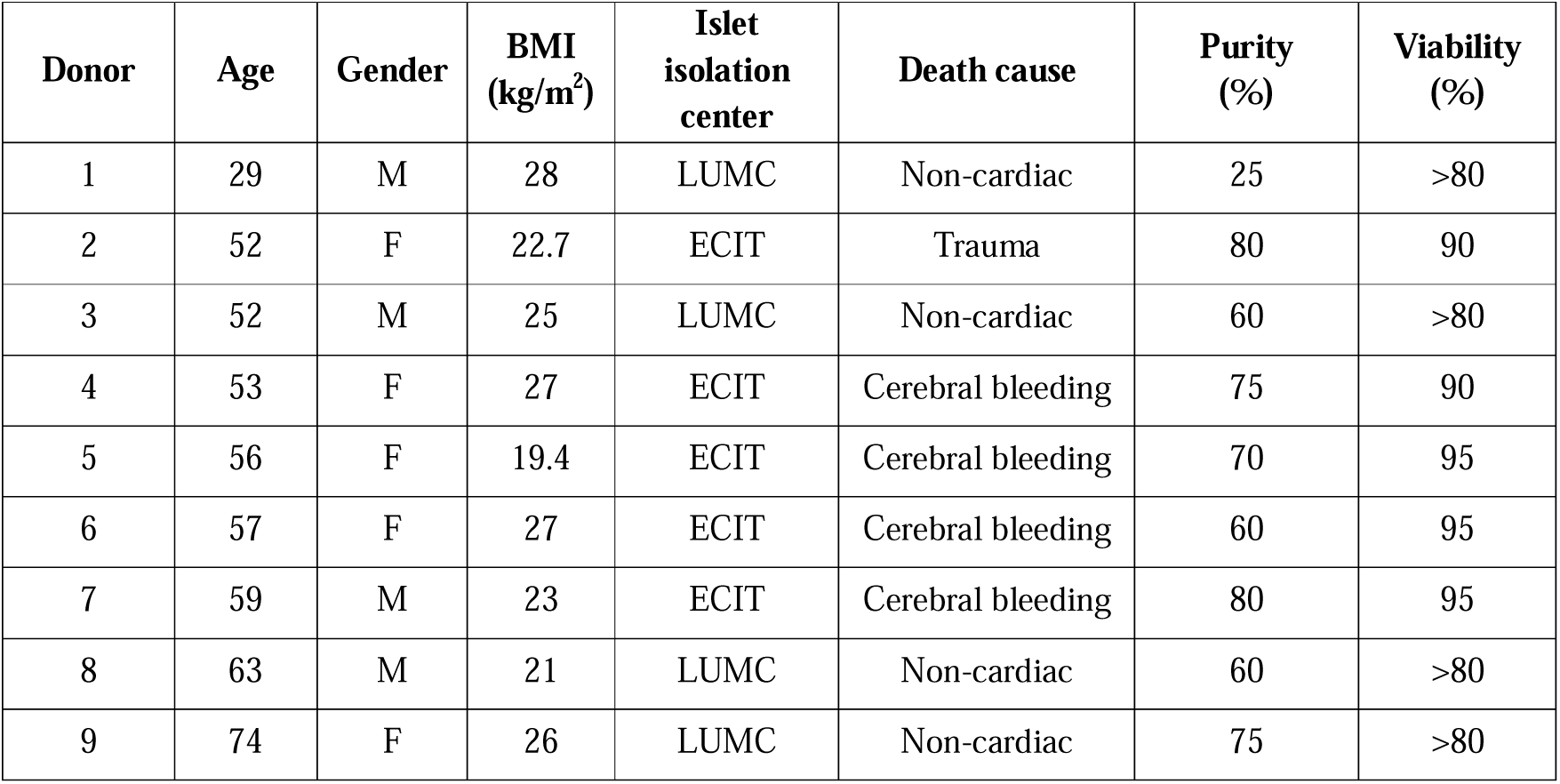
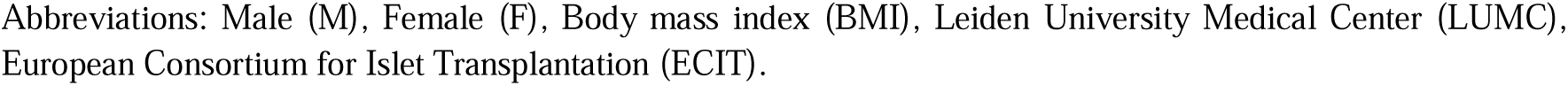
Characteristics of the human donors and pancreatic islets used in this study.

| Donor | Age | Gender | BMI<br>(kg/m<sup>2</sup>) | Islet<br>isolation<br>center | Death cause | Purity<br>(%) | Viability<br>(%) |
| --- | --- | --- | --- | --- | --- | --- | --- |
| 1 | 29 | M | 28 | LUMC | Non-cardiac | 25 | >80 |
| 2 | 52 | F | 22.7 | ECIT | Trauma | 80 | 90 |
| 3 | 52 | M | 25 | LUMC | Non-cardiac | 60 | >80 |
| 4 | 53 | F | 27 | ECIT | Cerebral bleeding | 75 | 90 |
| 5 | 56 | F | 19.4 | ECIT | Cerebral bleeding | 70 | 95 |
| 6 | 57 | F | 27 | ECIT | Cerebral bleeding | 60 | 95 |
| 7 | 59 | M | 23 | ECIT | Cerebral bleeding | 80 | 95 |
| 8 | 63 | M | 21 | LUMC | Non-cardiac | 60 | >80 |
| 9 | 74 | F | 26 | LUMC | Non-cardiac | 75 | >80 |

Abbreviations: Male (M), Female (F), Body mass index (BMI), Leiden University Medical Center (LUMC), European Consortium for Islet Transplantation (ECIT).

### 2.6. Rat pancreatic islet isolation

Rat islets were isolated from the same animals that served as donors for the r-prASC (described above). The isolation procedure followed a previously described protocol [20]. Briefly, the rat pancreas was distended by injecting ± 9 mL collagenase V solution (1 mg/mL; Sigma-Aldrich, Zwijndrecht, The Netherlands) in Hank’s Balanced Salt Solution (HBSS; Gibco, Thermo Fisher Scientific) via the pancreatic duct. Subsequently, the isolated islets were washed and purified using a Ficoll density gradient (Gradient stock solution; Corning Cellgro, Manassas, USA). Purified islets were cultured overnight in CMRL-M (+) at 37°C and 5% CO_2_. Acclimatized islets were used within 24 hours for secretome collection.

### 2.7. Secretome collection

For the co-culture conditions, h-prASC or r-prASC were seeded at 75.000 or 250.000 cells/well in non-adherent 6-well plates (Thermo Fisher Scientific) containing STD-M (+). These cell concentrations were determined using the ratios 1:300 and 1:1000 (islets to prASC). Plated cells were incubated overnight in a 5% CO_2_ incubator at 37 °C to allow adherence. After this period, adherent cells were carefully washed with phosphate-buffered saline (PBS) (Thermo Scientific), and 250 islets were added to each well. For the collection of secretome, the islets (250) or the co-cultures of islets and prASC (1:300 and 1:1000) were cultured in CMRL (−) – as this medium is the standard medium used for islets culturing – under the following conditions: normoxia (21% O_2_, 5% CO_2_), cytokines exposure (21% O_2_, 5% CO_2,_ 50 ng/mL IFN-γ, 21.5 ng/mL TNF-α, 0.25 ng/mL IL-1β), high glucose exposure (21% O_2_, 5% CO_2,_ 16.7 mM glucose), hypoxia (1% O_2_, 5% CO_2_), and hypoxia + high glucose exposure (1% O_2_, 5% CO_2,_ 16.7 mM glucose. After 72 h, the secretomes were harvested, pooled into three pools of three donors, and subjected to centrifugation (500 x g, 5 min) to remove undesired components. Secretomes were aliquoted and snap-frozen in liquid nitrogen and stored at −80°C until processing.

### 2.8. Discovery-based proteomics and data analysis

The protein composition of the various collected secretomes was determined using a discovery-based proteomics approach called label-free quantification [21]. Briefly, in-gel digestion was performed on 30 µL of the provided secretomes using trypsin (300 ng sequencing grade modified trypsin V5111; Promega, Leiden, The Netherlands) after first reducing them with 10 mmol/L dithiothreitol and then alkylating with 55 mmol/L iodoacetamide proteins [21]. Discovery mass spectrometric analyses were executed on a quadrupole orbitrap mass spectrometer equipped with a nano-electrospray ion source (Orbitrap Exploris 480; Thermo Scientific). Peptides were chromatographically separated using liquid chromatography (LC) on an Evosep system (Evosep One; Evosep, Odense, Denmark) with a nano-LC column (EV1137 Performance column 15 cm x 150 µm, 1.5 µm, Evosep). The LC-mass spectrometry (LC-MS) procedure used the equivalent of 1 µL starting injection material, and the peptides underwent separation using the 30SPD workflow (Evosep). The mass spectrometer was operated in positive ion mode and utilized the data-independent acquisition mode (DIA) with isolation windows of 16 m/z. The precursor mass range was set between 400 and 1000, and the FAIMS switched between CV-45V and −60V with three scheduled MS1 scans during each screening of the precursor mass range. The LC-MS raw data underwent processing using Spectronaut (version 17.1.221229; Biognosys Inc, Cambridge, United States), following the standard settings of the direct DIA workflow. Quantification was conducted on MS1, utilizing a human or rat SwissProt database (www.uniprot.org, containing 20,350 entries for human samples and 8,094 entries for rat samples). Local normalization was applied for quantification, and Q-value filtering was set to the classic setting without imputation. The final list of proteins identified in all conditions can be found in Supp. File 1 Table 1 (human) and Supp. File 2 Table 1 (rat).

### 2.9. Mass spectrometry *in silico* analysis

To gain insights into the potential functional changes occurring when islets and co-cultures are exposed to the various culturing strategies, the lists of proteins resulting from the secretomics studies were submitted to Metascape [22]. Metascape analyzed pathway enrichment clusters from Gene Ontology (GO) biological processes for each secretome. The top 100 biological processes from each secretome can be found in Supp. File 1 (human) and 2 (rat).

### 2.10. Quantification of paracrine factors

Within the secretomes, quantification of paracrine factors linked to angiogenesis (vascular endothelial growth factor (VEGF), platelet-derived growth factor AB (PDGF), and basic fibroblast growth factor (bFGF)), as well as the ECM constituent (collagen I alpha 1), and immunomodulatory proteins (IL-10 and IL-1α) was done using ELISA or Luminex techniques (Pinheiro-Machado, E., et al. (2024) Culturing conditions dictate the composition and pathways enrichment of human and rat perirenal adipose-derived stromal cells’ secretomes. Stem Cell Reviews and Reports. Advanced online publication.). Levels of rat and human bFGF, PDGF, VEGF (DuoSet ELISA; R&D Systems, Abingdon, UK), and rat collagen I alpha I (Novus Biologicals, Biotechne, Minneapolis, USA) were quantified through ELISA. Rat and human IL-10, IL-1α, and human collagen I alpha 1 were assessed utilizing magnetic Luminex® Assays (R&D systems; #LXSARM-03 / #LXSAHM-04). The secretome samples used for these assays consisted of three samples from each culturing group that were pooled (n = 3). These were then measured in triplicates following respective manufacturer protocols. For magnetic Luminex® Assays, plates were analyzed utilizing a Luminex 200 System, with data analysis conducted through Luminex xPONENT software. For ELISA kits (R&D Systems and Novus Biologicals), data acquisition was obtained using a microplate spectrophotometer (Epoch 2; BioTek, Winooski, USA) at wavelengths of 450 nm, with 540 nm correction applied.

### 2.11. Tube formation assay

To assess the proangiogenic capacity of the human secretomes, the Endothelial Cell Facility of the UMCG provided commercially obtained Human Umbilical Vein Endothelial Cells (HUVEC). The HUVEC were cultured in endothelial cell growth basal medium (EGM-2; Lonza) supplemented with EGM-2 MV SingleQuot Kit Supplements and Growth Factors (Lonza) at 37 °C with 5% CO_2_. Similarly, to assess the proangiogenic potential of rat secretomes, rat aortic endothelial cells (RAEC) were isolated according to a protocol previously outlined by Suh *et al.* [23]. For the assay, HUVEC (n = 3) and RAEC (n = 3) were seeded onto growth factor-reduced Matrigel (Corning, Amsterdam, The Netherlands) in a 96-well plate at a density of 35,000 cells per well. Secretomes were added, and cells were incubated for 18 hours at 37°C with 5% CO_2_. Post-incubation imaging was conducted using a Leica MZ7.5 microscope with a Leica IC90 E camera (Leica Microsystems B.V., Amsterdam, The Netherlands). HUVEC and RAEC cultivated in CMRL (−) served as control. The angiogenic potential was quantified based on the number of branching points counted by Fiji software [24]. Subsequently, the number of branching points was normalized to the control condition (CMRL (−)), set at 1.

### 2.12. Picrosirius Red staining

To assess the human secretomes’ capacity to induce collagen deposition, commercially acquired human fetal lung fibroblasts (FLF92, passage 28 - 30) were cultivated in STD-M (+). Comparatively, rat dermal fibroblasts (RDF) were utilized to assess the potential of the rat secretomes to stimulate collagen deposition. RDF were obtained through a serial explant technique as previously outlined by Nejaddehbashi *et al.* [25]. Picrosirius Red staining (Direct Red 80, Sigma-Aldrich) was performed to detect and quantify collagen deposition by FLF92 and RDF cells upon 72 h exposure to the different human or rat secretomes. The protocol of Xu *et al.* [26] was adapted for this purpose. The staining’s optical density (OD) was determined at 540 nm using a spectrophotometer (Epoch 2; BioTek, Winooski, USA). OD values were normalized to the total cell count (Countess 3 Cell Counter, ThermoFisher) following exposure to each secretome. Collagen content is expressed as OD per 10,000 cells, with ODs standardized to the control (CMRL (−)), set at a reference value of 1.

### 2.13. Two-way MLR followed by an antibody-mediated cell dependent cytotoxicity assay (CDC)

To elucidate the immunomodulatory capacity of both human and rat secretomes, we performed an adaptation of a previously described two-way MLR followed by an antibody-mediated CDC method [27, 28]. An overview of the procedure is depicted in Supplementary Fig 1. For human secretome, human peripheral blood mononuclear cells (PBMC) were used. For rat secretomes, rat splenocytes were used.

#### 2.13.1. Human peripheral blood mononuclear cells (PBMC)

To test the effect of different human prASC, isolated human peripheral blood mononuclear cells (PBMC) were used. To this end, buffy coats of healthy blood donors were obtained from the Sanquin Blood Bank (Groningen, The Netherlands). A Lymphoprep density gradient (Fresenius Kabin Norge AS, Oslo, Norway) and centrifugation (800 x g, 15 min, RT) were used to obtain the PBMC. Subsequent centrifugation to wash the cells were carried out at 300 x g (5 min, RT). The supernatant was discarded, and the interphase consisting of PBMC was transferred to new centrifuge tubes and washed three times with PBS. Before the final centrifugation, the PBMC were filtered through a 100 µm cell strainer (FALCON, Corning, Durham, USA). The viable cells were counted and immediately used for the MLR (counting and viability assessements were performed using a Countess 3 Cell Counter (ThermoFisher).

#### 2.13.2. Rat splenocytes

To investigate the immunomodulatory potential of the different rat prASC secretomes, rat splenocytes were used. To obtain those, spleens were collected from the same animals used for adipose tissue collection as previously described [29]. Briefly, the spleens were cut into small pieces and mechanically disrupted in ice-cold RPMI (+) (Gibco). Splenic red blood cells were eliminated by incubation with ice-cold ammonium chloride (4 mL, 10 min; UMCG Apotheek). Falcon tubes with cell strainer caps (35 μm; Corning) were used to remove cell clumps before the cells were counted and immediately plated for the MLR.

#### 2.13.3. Two-way MLR followed by CDC

The assay involved a two-way MLR, where stimulator and responder cells were co-cultured to generate alloantibodies. For the assay involving human cells, PBMC from mismatched human donors were co-cultured. In the case of rat cells, the co-culturing was conducted using splenocytes obtained from two different rat strains, with Sprague Dawley rats serving as stimulators/resting and Wistar rats as responders. Briefly, stimulator cells (0.5 × 10^6^ cells per well) were treated with mitomycin C (50 μg/mL; Sigma-Aldrich; Merck Millipore) for 30 min at 37 °C in a 24-well plate. After treatment, mitomycin C-treated stimulator cells were washed three times with RPMI (+) and centrifuged at 300 x g for 5 min at RT. Subsequently, these cells were then resuspended in DMEM (+) or the different prASC secretomes and placed into a 24-well plate (n = 10 for h-prASC secretomes and n = 7 for r-prASC secretomes). Responder cells (0.5 × 10^6^ cells; also resuspended in DMEM (+) or the different prASC secretomes) were added to each well containing stimulator cells. Together, stimulator and responder cells were cultured for seven days at 37 °C and 5% CO_2_. During this period, no media change was performed.

In parallel, resting splenocytes or PBMC (from the same donor as the stimulator cells; 10^6^ cells) were washed twice with RPMI (+), centrifuged at 300 x g for 5 min at RT, resuspended in RPMI (+) and seeded onto a 6-well plate. Resting splenocytes were also incubated for seven days at 37 °C, and 5% CO_2_, with no media changes in this period. Upon completion of the 7-day incubation, the supernatants resulting from the one-way-MLR (MLR supernatants, containing alloantibodies resulting from the exposure or not of the different secretomes) were collected (450 uL). Additionally, the number and viability of the resting cells was determined (Countess 3 Cell Counter (ThermoFisher)).

The second part of the assay evaluated humoral alloimmunity using the antibody-mediated complement-derived cytotoxicity assay (CDC assay). For it, we cultured the previously counted resting cells in the different MLR supernatants. Briefly, resting cells (0.5 × 10^5^) were resuspended in the respective MLR supernatants (50 uL) and seeded onto a 96-well plate. STD-M (+) served as control. Resting splenocytes were cultured with the supernatants for 30 min at RT. After this, low toxicity rabbit complement (Sanbio B.V., Cedarlane, Uden, The Netherlands) was added to each well and incubated for 2 h, 37 °C, and 5% CO_2_.

To evaluate the resting splenocytes’ survival upon exposure to the different MLR supernatants (with or without the presence of secretomes) or controls (STD-M (+)), a WST-1 assay (Sigma-Aldrich) was performed following the manufacturer’s instructions. The capacity of each secretome to mitigate alloimmunity was defined as the % of human PBMC or rat splenocytes that survived upon exposure to the MLR supernatants. Cell survival was normalized to the control (STD-M (+); set at 100).

### 2.14. Statistical analysis

Data are expressed as mean ± standard deviation (SD). Statistical analysis was carried out in GraphPad Prism (version 9.2.0; GraphPad Software, Inc., La Jolla). For all *in vitro* functional assays, the effect of the different conditions, co-culturing, and the interaction between these two was analyzed by a two-way ANOVA (TWA). To do so, the normality of the data was first tested using the Kolmogorov–Smirnov test. If data were not normally distributed, the data was log-transformed before performing the TWA. Šídák’s multiple comparison test was used as post-testing. To obtain insights into the effect of the secretome from islets cultured alone, the islet secretomes from all conditions were compared to the islet normoxia-derived secretome. To obtain insights into the effect of the secretome from co-cultured islets, the co-cultured islets normoxia-derived secretome was also used for comparison. Moreover, to compare the effect of the different co-cultured islet secretomes, we compared all conditions to each other. Data were considered significantly different if p < 0.05.

## 3. Results

### 3.1. The pancreatic islets secretome after different culturing conditions

The secretomes of *in vitro*-cultured human and rat pancreatic islets were characterized to assess alterations in the islets’ secretion profile when encountering diverse stress conditions that mimic the situation during isolation and after PIT. Leveraging secretomics data, which includes a comprehensive list of identified proteins for each condition, Metascape was used to discern the top 100 most enriched pathways within each condition. All pathway enrichment analyses were conducted using the complete list of identified proteins from each condition’s secretome. Below the five most enriched pathways are discussed for each secretome. Furthermore, additional pathways of interest are highlighted, specifically those associated with angiogenesis, ECM, and immune responses, since these pathways are crucial for functional islet survival and PIT success. The baseline secretion signature was established using the normoxia-derived secretome for each species.

#### 3.1.1. The human islet secretome

##### 3.1.1.1. Proteomics of human normoxia-derived secretome

In the normoxia-derived secretome of human islets, 367 proteins were identified (Fig. 2A). The pathway enrichment analysis specific to this secretome revealed the following top five enriched pathways associated with these proteins: regulation of body fluids, wound healing, hemostasis, cellular response to toxic substances, and regulation of proteolysis (Table 2; Supp. File 1 Table 3). Additional pathways of interest in this secretome, which are still within the top 100, encompassed cellular response to IL-7, chronic inflammatory response, collagen metabolic process, regulation of humoral immune response, and blood vessel development (Supp. File 1 Table 3).

**Fig. 2.**
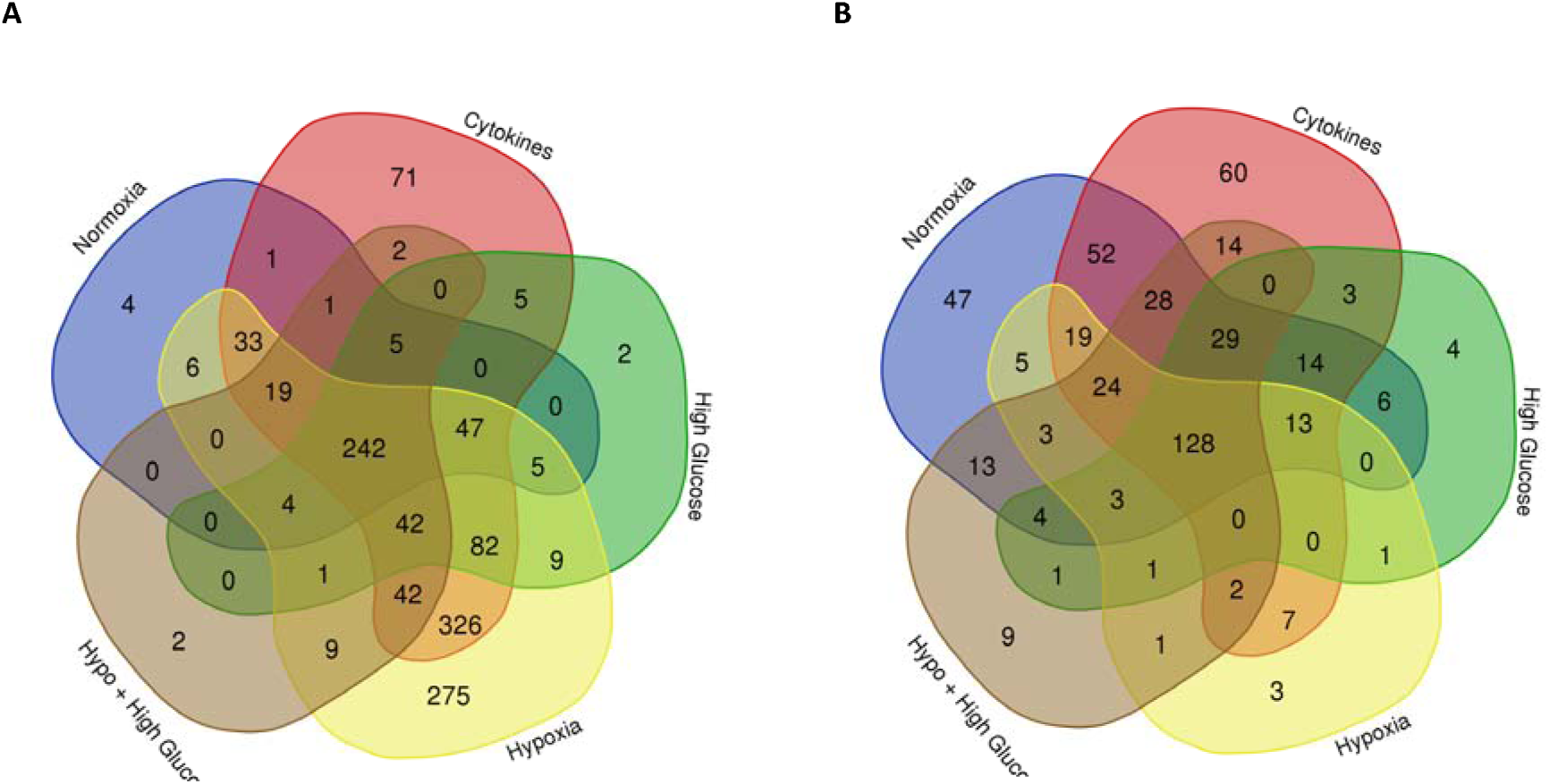
The human and rat islet secretome. Global overview of human (A) and rat (B) pancreatic islet secretomes resulting from various *in-vitro* culturing conditions.

**Table 2.** Human islets secretome.

|  | Normoxia-derived secretome |  | Cytokine-derived secretome |  | High glucose-derived secretome |  | Hypoxia-derived secretome |  | Hypoxia + High glucose-derived secretome |  |
| --- | --- | --- | --- | --- | --- | --- | --- | --- | --- | --- |
|  | Pathway | LogP | Pathway | LogP | Pathway | LogP | Pathway | LogP | Pathway | LogP |
| **Most enriched** | regulation of body fluid levels | -30,86 | generation of precursor metabolites and energy | -50,24 | regulation of body fluid levels | -33,69 | generation of precursor metabolites and energy | -57,54 | regulation of body fluid levels | -26,39 |
|  | wound healing | -29,20 | protein maturation | -33,89 | hemostasis | -32,16 | peptide metabolic process | -40,29 | wound healing | -25,79 |
|  | hemostasis | -27,30 | response to wounding | -33,25 | wound healing | -31,24 | cellular respiration | -38,34 | hemostasis | -21,02 |
|  | cellular response to toxic substance | -22,05 | carbohydrate metabolic process | -31,19 | generation of precursor metabolites and energy | -22,49 | protein maturation | -35,86 | regulation of proteolysis | -20,57 |
|  | regulation of proteolysis | -20,62 | monocarboxylic acid metabolic process | -29,97 | protein maturation | -21,25 | amide biosynthetic process | -35,83 | protein maturation | -19,97 |

**Table 3.** Rat islets secretome.

|  | Normoxia-derived secretome |  | Cytokine-derived secretome |  | High glucose-derived secretome |  | Hypoxia-derived secretome |  | Hypoxia + High glucose-derived secretome |  |
| --- | --- | --- | --- | --- | --- | --- | --- | --- | --- | --- |
|  | Pathway | LogP | Pathway | LogP | Pathway | LogP | Pathway | LogP | Pathway | LogP |
| **Most enriched** | response to xenobiotic stimulus | -16,24 | response to xenobiotic stimulus | -16,81 | protein maturation | -14,24 | protein maturation | -12,43 | protein folding | -12,58 |
|  | response to metal ion | -13,90 | response to metal ion | -13,13 | protein folding | -11,33 | hemostasis | -12,26 | protein maturation | -11,53 |
|  | protein maturation | -12,96 | protein catabolic process | -11,99 | hemostasis | -11,14 | response to wounding | -11,58 | response to metal ion | -10,34 |
|  | response to wounding | -12,25 | generation of precursor metabolites and energy | -11,53 | response to metal ion | -10,66 | regulation of body fluid levels | -11,50 | monosaccharide biosynthetic process | -9,76 |
|  | protein catabolic process | -12,08 | monosaccharide metabolic process | -10,97 | wound healing | -10,23 | hydrogen peroxide metabolic process | -11,31 | response to xenobiotic stimulus | -8,78 |

##### 3.1.1.2. Proteomics of human cytokines-derived secretome

Exposure of human islets to cytokines, mimicking an inflammatory environment, resulted in the identification of 918 proteins (Fig. 2A). Among these, 570 (62.1%) were newly identified (when compared to the normoxia-derived secretome). The cytokines-derived secretome was enriched in the following pathways (top 5 pathways): generation of precursor metabolites and energy, protein maturation, response to wounding, carbohydrate metabolic processes, and monocarboxylic acid metabolic processes (Table 2; Supp. File 1 Table 4). Additionally, pathways of interest in this secretome encompassed innate immune response, cellular response to cytokine stimulus, and blood vessel development (Supp. File 1 Table 4).

**Table 4.**
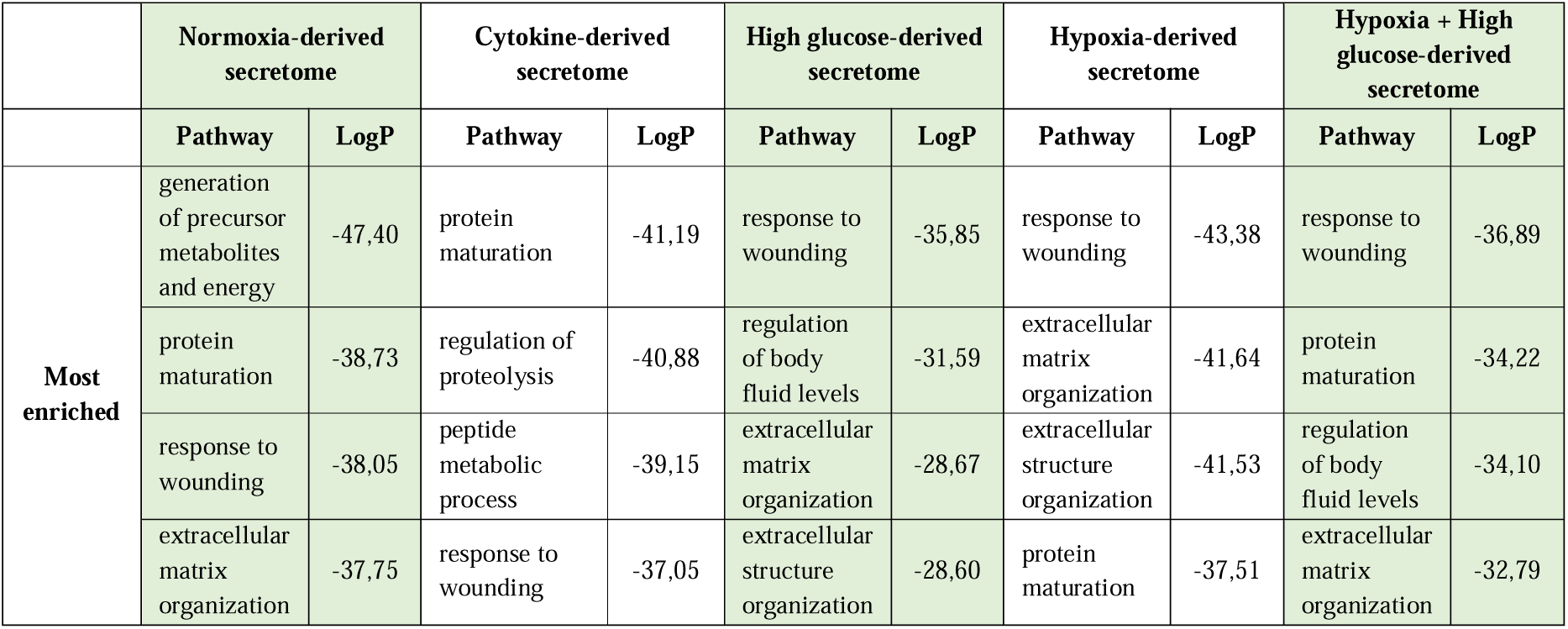
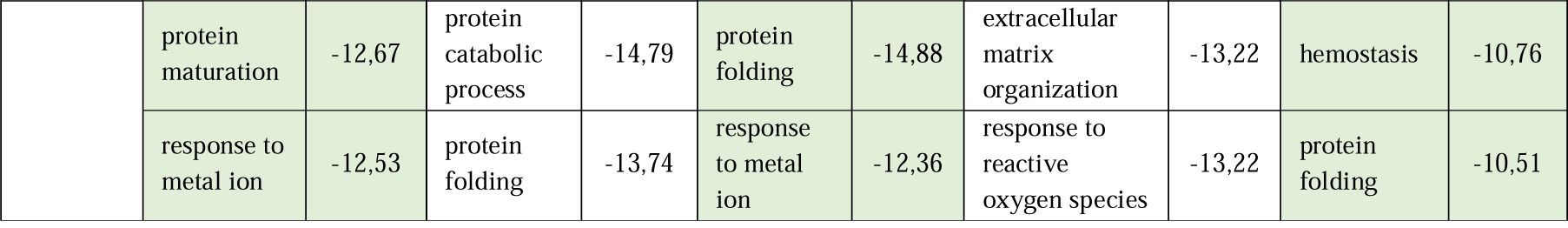
Human co-culture secretome (1:1000).

##### 3.1.1.3. Proteomics of human high glucose-derived secretome

Exposure to high glucose, simulating a hyperglycemic environment, identified 444 proteins in the islet secretome (Fig. 2A). Of these, 140 (31.5%) were newly secreted compared to the normoxia-derived secretome. This secretome was enriched in pathways of regulating body fluid levels, hemostasis, wound healing, generating precursor metabolites and energy, and protein maturation (Table 2; Supp. File 1 Table 5). Other enriched pathways included supramolecular fiber organization, innate immune response, ECM organization, regulating complement activation, endothelial cell development, angiogenesis, and more (Supp. File 1 Table 5).

**Table 5.** Rat co-culture secretome (1:1000).

|  | Normoxia-derived secretome |  | Cytokine-derived secretome |  | High glucose-derived secretome |  | Hypoxia-derived secretome |  | Hypoxia + High glucose-derived secretome |  |
| --- | --- | --- | --- | --- | --- | --- | --- | --- | --- | --- |
|  | Pathway | LogP | Pathway | LogP | Pathway | LogP | Pathway | LogP | Pathway | LogP |
| **Most enriched** | response to xenobiotic stimulus | -17,75 | response to xenobiotic stimulus | -19,45 | response to xenobiotic stimulus | -18,23 | response to xenobiotic stimulus | -18,45 | response to wounding | -13,81 |
|  | response to wounding | -13,34 | response to metal ion | -15,02 | response to wounding | -15,82 | supramolecular fiber organization | -15,48 | protein maturation | -12,27 |
|  | protein folding | -12,75 | protein maturation | -15,00 | protein maturation | -15,08 | protein maturation | -14,51 | response to xenobiotic stimulus | -12,13 |
|  | protein maturation | -12,67 | protein catabolic process | -14,79 | protein folding | -14,88 | extracellular matrix organization | -13,22 | hemostasis | -10,76 |
|  | response to metal ion | -12,53 | protein folding | -13,74 | response to metal ion | -12,36 | response to reactive oxygen species | -13,22 | protein folding | -10,51 |

##### 3.1.1.4. Proteomics of human hypoxia-derived secretome

The exposure of human islets to hypoxia, simulating the lack of vascularization post islet transplantation, resulted in a secretome with 1142 proteins (Fig. 2A), in which 786 (68.8%) were newly secreted when compared to the normoxia-derived secretome. The human islet hypoxia-derived secretome was enriched in pathways such as generation of precursor metabolites and energy, peptide metabolic process, cellular respiration, protein maturation, and amide biosynthetic process (Table 2; Supp. File 1 Table 6). Also, present in this secretome were pathways of supramolecular fiber organization, ECM organization, innate and humoral immune system, and blood vessel development, among others (Supp. File 1 Table 6).

##### 3.1.1.5. Proteomics of human hypoxia + high glucose-derived secretome

Exposing islets to a combination of hypoxia and high glucose not only replicates physiological conditions in individuals with diabetes (hyperglycemia and insufficient oxygen supply) but also simulates a post-transplantation environment in an islet transplantation context. In the islet secretome under hypoxia and high glucose conditions, 369 proteins were identified, and 98 of those were newly secreted when compared to the normoxia-derived secretome (26.5%) (Fig. 2A). Enriched pathways included regulation of body fluid levels, wound healing, hemostasis, regulation of proteolysis, and protein maturation (Table 2; Supp. File 1 Table 7). Other pathways of interest also identified in this secretome included supramolecular fiber organization, humoral immune response, ECM organization, blood vessel development, blood vessel endothelial cell migration, and others (Supp. File 1 Table 7).

#### 3.1.2. The rat islet secretome

##### 3.1.2.1. Proteomics of rat normoxia-derived secretome

In the rat islet normoxia-derived secretome, 388 proteins were identified (Fig. 2B). The pathway enrichment analysis indicated their primary association with processes related to response to xenobiotic stimulus, response to metal ion, protein maturation, response to wounding, and protein catabolic process (Table 3; Supp. File 2 Table 3). Other pathways of interest in this secretome include response to interleukin-1 (IL-1) and IL-7, regulation of oxygen species metabolic process, and blood vessel development (Supp. File 2 Table 3).

##### 3.1.2.2. Proteomics of rat cytokines-derived secretome

When exposed to cytokines, the rat islet secretome contained 393 proteins (Fig. 2B). From those, 86 (21.9%) new proteins emerged compared to the normoxia-derived secretome. The presence of these proteins has enriched this secretome in pathways of response to xenobiotic stimulus, response to metal ion, protein catabolic process, generation of precursor metabolites and energy, and monosaccharide metabolic process (Table 3; Supp. File 2 Table 4). Other pathways of interest enriched in the rat islet cytokines-derived secretome includes collagen fibril organization, supramolecular fiber organization, insulin metabolic process, cellular response to IL-1 and IL-7, and complement activation (Supp. File 2 Table 4).

##### 3.1.2.3. Proteomics of rat high glucose-derived secretome

Exposing rat islets to high glucose yielded a secretome containing 207 proteins (Fig. 2B). Only 10 (4.83%) new proteins were identified as newly secreted compared to the normoxia-derived secretome. Pathways such as protein maturation, protein folding, hemostasis, response to metal ion, and wound healing composed this secretome’s top 5 most enriched processes (Table 3; Supp. File 2 Table 5). Other enriched pathways of interest included ECM organization, insulin metabolic process, supramolecular fiber organization, vasculature development, collagen metabolic process, and complement activation (Supp. File 2 Table 5).

##### 3.1.2.4. Proteomics of rat hypoxia-derived secretome

The secretome derived from rat islets exposed to hypoxia resulted in 220 proteins (Fig. 2B). From those, only 15 (6.82%) new proteins were identified when compared to the normoxia-derived secretome. The top 5 pathways enriched included protein maturation, hemostasis, response to wounding, regulation of body fluid levels, and hydrogen peroxide metabolic process (Table 3; Supp. File 2 Table 6). Other pathways of interest identified in the rat islet hypoxia-derived secretome included supramolecular fiber organization, and cellular response to IL-7, among others (Supp. File 2 Table 6).

##### 3.1.2.5. Proteomics of rat hypoxia + high glucose-derived secretome

Finally, in the rat islet hypoxia + high glucose-derived secretome, 260 proteins were identified, and 28 (10.8%) from those were newly secreted when compared to the normoxia-derived secretome (Fig. 2B). Pathway enrichment revealed protein folding, protein maturation, response to metal ion, monosaccharide biosynthetic process, and response to xenobiotic stimulus as the top 5 highly enriched pathways in this secretome (Table 3; Supp. File 2 Table 7). Other pathways of interest included cellular response to IL-7, collagen fibril organization, complement activation, regulation of cellular response to stress, and positive regulation of angiogenesis, among others (Supp. File 2 Table 7).

### 3.2. The impact of ASC and islet co-culturing on the secretome under different culturing conditions

ASC have shown to improve pancreatic islet function and transplantation after isolation and PIT [6, 13, 15]. To investigate if changes in the secretome profile might be involved in this beneficial effect, the secretome of human and rat islets co-cultured with ASC under various stress conditions was investigated. The secretomes of two different ratios of islets vs. ASC (1:300 and 1:1000) were collected for human (Supp. File 1 Table 8) and rat (Supp. File 2 Table 8) co-cultures. Using the identified proteins for each condition, a pathway enrichment analysis was conducted, focusing on GO biological processes to understand the functional profile of these secretomes. Prior to this, the influence of ASC proportion (1:300 or 1:1000) on the secretome of the co-culture was assessed through a comparative analysis of the total proteins in the normoxia-derived secretome of islets cultured alone or with ASC in the ratios 1:300 and 1:1000 (Fig. 3). This served as an indicator to discern whether a lower islet to ASC ratio could already exert a significant influence on secretome composition.

**Fig. 3.**
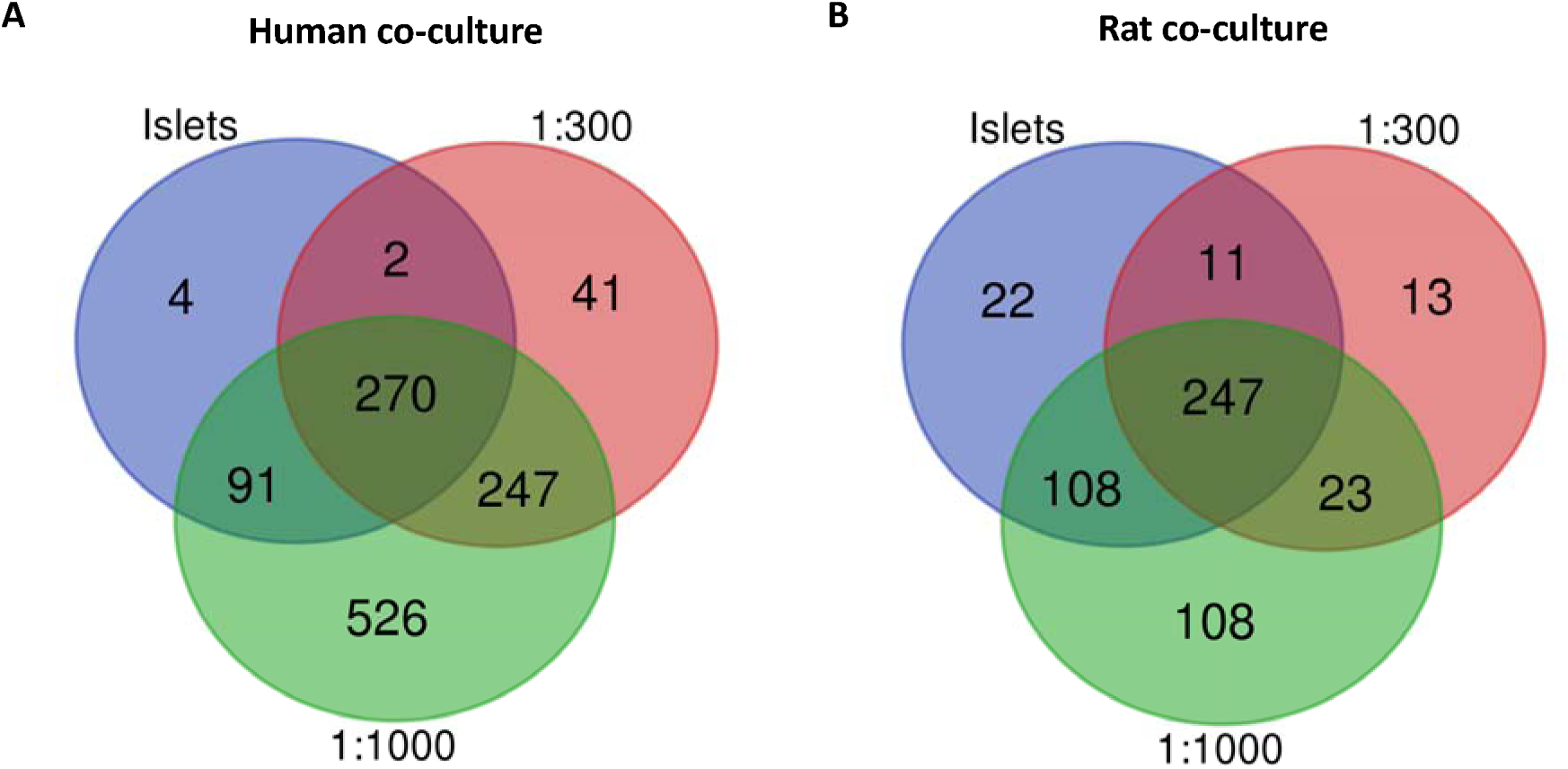
Modulation of the human and rat islet secretome composition upon co-culturing with ASC. Global overview of human (A) and rat (B) islets co-cultured or not with adipose-derived stromal cells (ASC; either in a 1:300 or a 1:1000 islet to ASC ratio).

For human islets, the 1:300 co-culture normoxia-derived secretome displayed 288 new proteins when compared to the normoxia-derived human islet secretome (constituting 78.5% of the total identified in this secretome; Fig. 3A), while the 1:1000 co-culture secretome exhibited 773 newly identified proteins (68.17% of the total identified in this secretome; see Fig. 3A). For rat islets, the 1:300 co-culture secretome revealed 36 new proteins when compared to the rat islet normoxia-derived secretome (12.2% of the total identified, Fig. 3B). In comparison, the 1:1000 co-culture secretome displayed 131 proteins (27% of the total identified). Notably, a higher abundance of newly identified proteins was observed in the 1:1000 co-culture secretome compared to the 1:300, indicating that more pronounced changes could be identified in the functional profile of this co-culture. Subsequently, our analyses focused on exploring the pathway enrichment of secretomes of 1:1000 co-cultures under both standard and stress conditions of both species.

#### 3.2.1. The human islet and ASC co-culture secretome

##### 3.2.1.1. Proteomics of human normoxia-derived co-culture secretome

In the human co-culture normoxia-derived secretome, 1134 proteins were identified (Fig. 4A; Supp. File 1, Table 9). Comparing this secretome to the one from islets alone exposed to normoxia, 773 new proteins were found, constituting 68.1% of the total proteins identified (Fig. 4A). Enriched pathways were generation of precursor metabolites and energy, protein maturation, response to wounding, ECM organization, regulation of proteolysis, among others (Table 4; Supp. File 1 Table 9). Additionally, other pathways of interest were identified within this secretome. Those included the top 5 enriched pathways previously described in the normoxia-derived secretome from islets alone (Table 2), as well as pathways related to innate immune responses, humoral immune responses mediated by circulating immunoglobulins, complement activation, and collagen fibril organization (Supp. File 1, Table 9).

**Fig. 4.**
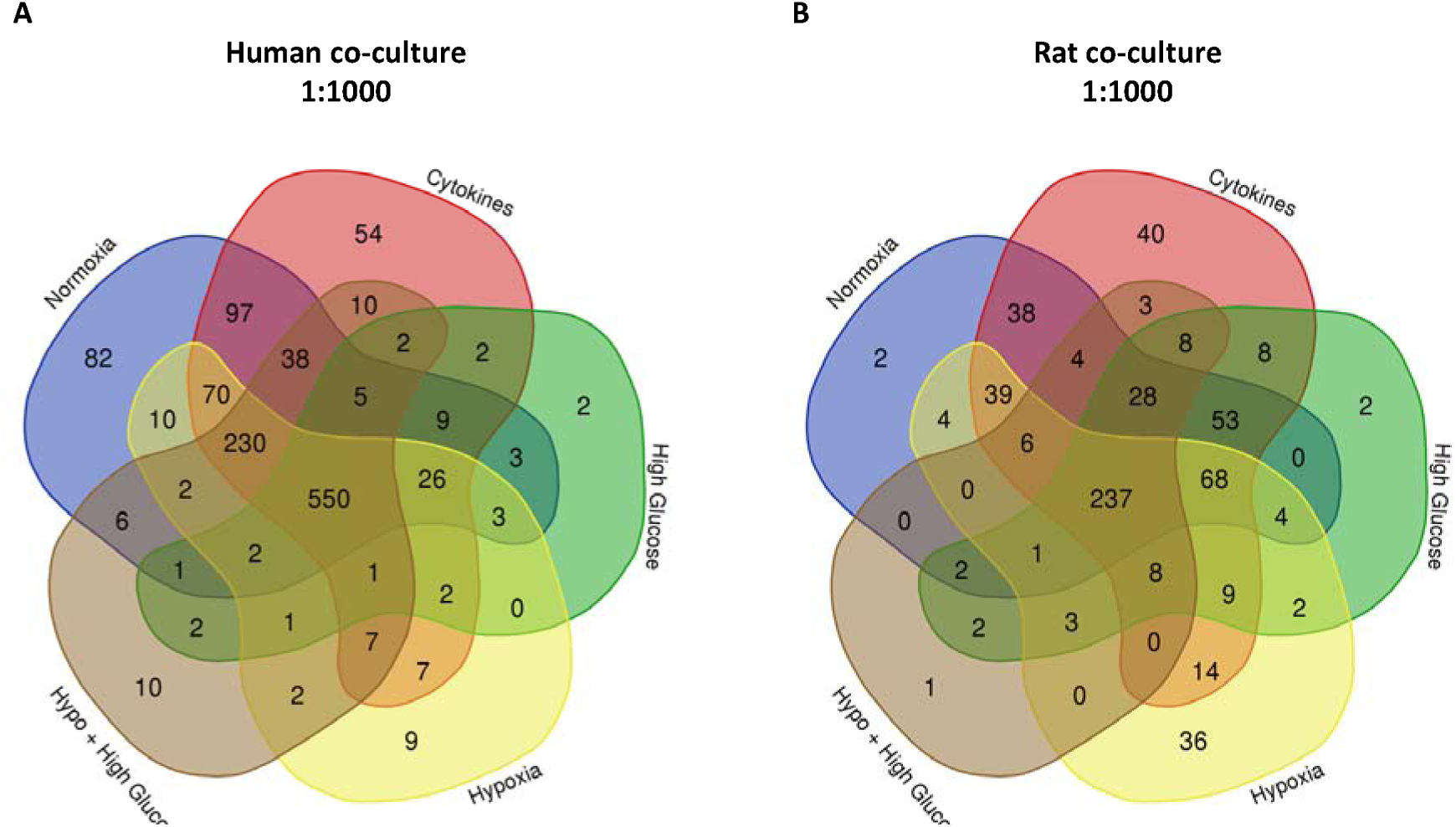
The human and rat co-cultured pancreatic islet secretome. Global protein overview of secretomes derived from human (A) and rat (B) islets co-cultured with adipose-derived stromal cells in a ratio 1:1000 under various in vitro culture conditions.

##### 3.2.1.2. Proteomics of human cytokines-derived co-culture secretome

In the human co-culture secretome derived under cytokines exposure, 1110 proteins were detected (Fig. 4A; Supp. File 1, Table 10). Among them, 246 new proteins were found when compared to the normoxia-derived co-culture secretome, constituting 22.2% of the total (Fig. 4A). This secretome demonstrated enrichment in pathways related to protein maturation, regulation of proteolysis, peptide metabolic process, response to wounding, and ECM organization (Table 4; Supp. File 1 Table 10). Other pathways of interest included the top 5 enriched pathways previously described in the cytokines-derived secretome from islets alone (Table 2) and pathways related to innate and humoral immune responses, collagen fibril organization, and immunoglobulin-mediated immune responses (Supp. File 1, Table 10).

##### 3.2.1.3. Proteomics of human high glucose-derived co-culture secretome

Under high glucose human co-culture conditions, 611 proteins were found (Fig. 4A; Supp. File 1, Table 11), with 246 new proteins when compared to the normoxia-derived co-culture secretome, making up 40.3% of the total (Fig. 4A). This secretome exhibited enrichment in pathways related to response to wounding, regulation of body fluid levels, ECM organization, extracellular structure organization, and negative regulation of peptidase activity (Table 4; Supp. File 1 Table 11). Furthermore, additional pathways of interest included the top 5 enriched pathways observed in the high glucose-derived secretome from islets alone (Table 2), and also pathways of supramolecular fiber organization, innate immune responses, collagen metabolic processes, developments in the vasculature, blood vessel formation, and morphogenesis of blood vessels (Supp. File 1, Table 11).

##### 3.2.1.4. Proteomics of human hypoxia-derived co-culture secretome

In hypoxic human co-cultures, we detected 922 proteins (Fig. 4A; Supp. File 1 Table 12), and 153 (16.6%) of them were newly secreted when compared to the normoxia-derived co-culture secretome (Fig. 4A). This secretome displayed enrichment in pathways linked to the response to wounding, the organization of the ECM and its structural components, protein maturation, and the regulation of body fluid levels (Table 4; Supp. File 1 Table 12). Furthermore, additional pathways of interest included those that corresponded with the top 5 enriched pathways previously observed in the hypoxia-derived secretome from islets alone (Table 2), as well as humoral immune responses, the organization of collagen fibrils, developments in blood vessel formation, and humoral immune responses mediated by circulating immunoglobulins (Supp. File 1 Table 12).

##### 3.2.1.5. Proteomics of human hypoxia + high glucose-derived co-culture secretome

Finally, under both hypoxia and high glucose conditions, 869 human proteins were identified (Fig. 4A; Supp. File 1 Table 13) and 544 (62.6%) proteins were newly secreted when compared to the normoxia-derived co-culture secretome (Fig. 4A). This secretome showed enrichment in pathways associated with the response to wounding, protein maturation, the regulation of body fluid levels, extracellular matrix organization, and the regulation of proteolysis (Table 5; Supp. File 1 Table 13). Furthermore, additional enriched pathways of interest included the top 5 previously observed in the hypoxia + high glucose-derived secretome from islets alone (Table 2), supramolecular fiber organization, and innate immune responses (Supp. File 1 Table 13).

#### 3.2.2. The rat islet and ASC co-culture secretome

##### 3.2.2.1. Proteomics of rat normoxia-derived co-culture secretome

When cultured in standard normoxic conditions, the co-culture between rat islets and rat ASC resulted in a secretome containing 486 proteins (Fig. 4B). Comparing this secretome to the one from islets alone exposed to normoxia, 131 new proteins were found, constituting 26.9% of the total proteins identified (Fig. 4B). The pathway analysis unveiled significant enrichment processes such as response to xenobiotic stimulus, response to wounding, protein folding, protein maturation, and response to metal ion (Table 5; Supp. File 2 Table 9). Noteworthy connections were observed with the top 5 enriched pathways previously identified in the normoxia-derived secretome from islets alone (Table 3; Supp. File 2 Table 3). Other pathways of interest, such as supramolecular fiber organization, generation of precursor metabolites and energy, and ECM, were identified (Supp. File 2 Table 9). Surprisingly, the top 100 pathways in this secretome did not include any pathways directly associated with the immune system or blood vessel development (Supp. File 2 Table 9).

##### 3.2.2.2. Proteomics of rat cytokines-derived co-culture secretome

The rat co-culture cytokine-derived secretome contained 563 proteins (Fig. 4B; Supp. File 2 Table 10). Among these, 187 proteins (33.2%) were newly secreted when compared to the normoxia-derived co-culture secretome (Supp. File 2 Table 10). Enriched pathways in this mixture involved response to xenobiotic stimulus, response to metal ion, protein maturation, protein catabolic process, protein folding (Table 5; Supp. File 2 Table 10). Other pathways of interest included those already described in the top 5 enriched pathways of the cytokines-derived islet secretome (Table 3; Supp. File 2 Table 4), as well as supramolecular fiber organization (Supp. File 2 Table 10). Interestingly, no immune system, blood vessel development, or additional ECM-directly associated pathways were identified within the top 100 in this secretome (Supp. File 2 Table 10).

##### 3.2.2.3. Proteomics of rat high glucose-derived co-culture secretome

The investigation into the rat co-culture high glucose-derived secretome revealed the identification of 435 proteins (Fig. 4B; Supp. File 2 Table 11). From those, 249 proteins (57.2% of the total) were newly secreted when compared to the normoxia-derived co-culture secretome (Supp. File 2 Table 11). The pathway analysis unveiled pathways such as response to xenobiotic stimulus, response to wounding, protein maturation, protein folding, and response to metal ion, as highly enriched (Table 5; Supp. File 2 Table 11). Other pathways of interest described within the top 100 included not only the previously described top 5 most enriched of the rat islet only high glucose-derived secretome (Table 3; Supp. File 2 Table 5) but also pathways of supramolecular fiber organization, regulation of endothelial cell migration, response to interleukin-1 and interleukin-6, and response to insulin, among others (Supp. File 2 Table 11).

##### 3.2.2.4. Proteomics of rat hypoxia-derived co-culture secretome

Rat co-cultures exposed to hypoxia secreted 431 proteins (Fig. 4B; Supp. File 1 Table 12). Among these, 249 (57.8%) were identified as newly secreted when compared to the normoxia-derived co-culture secretome (Supp. File 1 Table 12). The pathway analysis revealed significant enrichment in response to xenobiotic stimulus, supramolecular fiber organization, protein maturation, ECM organization, and response to reactive oxygen species (Table 6; Supp. File 1 Table 12). Other pathways of interest identified in this secretome included those already described as the top 5 most enriched in the rat islet only hypoxia-derived secretome (Table 3; Supp. File 2 Table 6). Additionally, positive regulation of endothelial cell migration, collagen fibril organization, and others were identified as enriched pathways (Supp. File 1 Table 12). Notably, no pathways directly associated with immune system processes were included in this secretome’s top 100 most enriched pathways.

##### 3.2.2.5. Proteomics of rat hypoxia + high glucose-derived co-culture secretome

Finally, in the secretome of rat co-cultures exposed to a combination of hypoxia and high glucose, we identified 303 proteins (Fig. 4B; Supp. File 1 Table 13). From these, 118 (40% of the total) were newly secreted when compared to the normoxia-derived co-culture secretome (Supp. File 1 Table 13). The pathway enrichment analysis revealed significant involvement of these proteins in various processes, including the response to wounding, protein maturation, response to xenobiotic stimulus, hemostasis, protein folding (Table 5; Supp. File 1 Table 13). Other pathways of interest were enriched, including the top 5 most enriched pathways observed in the rat islets only hypoxia + high glucose-derived secretome (Table 3; Supp. File 2 Table 7), as well as ECM organization, regeneration, collagen metabolic process, negative regulation of intrinsic apoptotic signaling pathway, and positive regulation of vascular associated smooth muscle cell proliferation (Supp. File 1 Table 13).

### 3.3. Identification of key factors in the secretome of islets cultured with or without ASC

ELISA and Luminex techniques were applied to validate the presence of key proteins associated with important biological processes essential for promoting optimal PIT outcomes. Specifically, we focused on VEGF, PDGF, and b-FGF, primarily linked to angiogenesis; Collagen I alpha I, a major ECM component; and IL-1α and IL-10, representing pro-and anti-inflammatory factors. This validation was performed across different secretomes, comparing islets cultured alone versus those co-cultured with ASC.

#### 3.3.1. The concentration of key proteins in secretomes from human islets with or without ASC

Figure 5 illustrates the analysis of key proteins involved in the pathways of interest within the secretome of islets cultured alone or with ASC. For VEGF (Fig. 5A), the TWA revealed significant effects of the culturing conditions (p<0.0001), the presence of ASC (p<0.0001), and their interaction (p<0.0001). Comparing the secretomes derived from islets cultured alone to their baseline (normoxia-derived) secretome, we observed a significant increase in VEGF in both hypoxia (Šídák’s post-test, p<0.0001) and hypoxia + high glucose-derived secretomes (Šídák’s post-test, p<0.0001). Similarly, a significant increase in VEGF levels was noted when comparing the secretomes from islets co-cultured with ASC to their baseline (normoxia-derived secretome) – both hypoxia (Šídák’s post-test, p<0.0001) and hypoxia + high glucose-derived secretomes (Šídák’s post-test, p<0.0001) displayed elevated VEGF levels. Further analysis of secretomes from islets cultured with ASC compared to those from islets cultured alone showed a significant increase in VEGF in the hypoxia-derived (Šídák’s post-test, p<0.0001) and hypoxia + high glucose-derived co-culture secretome (Šídák’s post-test, p<0.0001).

**Fig. 5.**
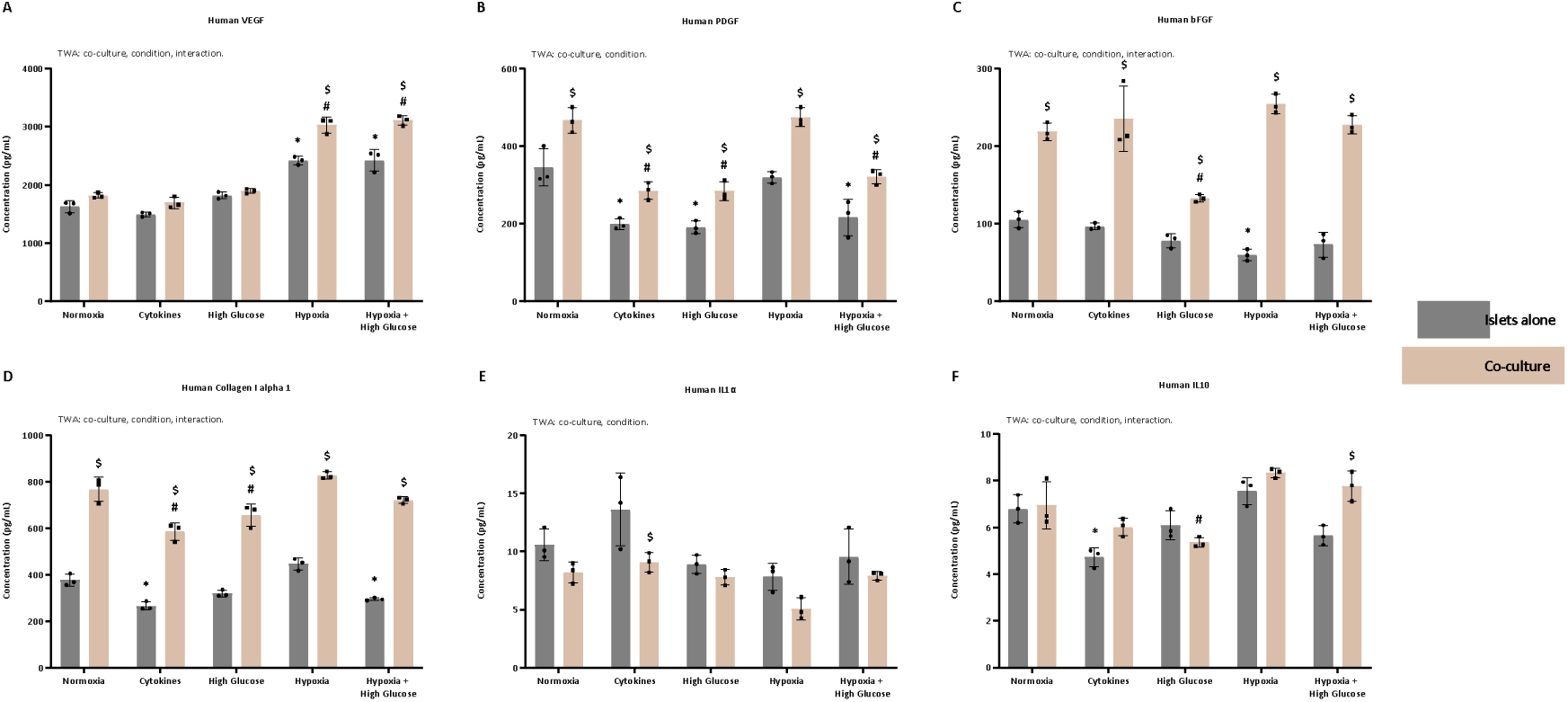
Modulation of key paracrine factors involved in pathways of interest in the human secretome upon co-culturing. Secretion of (A) vascular endothelial growth factor (VEGF), (B) platelet-derived growth factor (PDGF), (C) basic fibroblast growth factor (b-FGF), (D) collagen I alpha I, (E) interleukin 1 alpha (IL-1α), and (F) IL-10 after 72 hours co-culturing in standard conditions (21% O2). Data represent mean values ±0standard deviation of 3 pooled samples measured in triplicate. Two-way ANOVA (TWA) followed by Šídák’s multiple comparison test, * p < 0.05 versus Normoxia islets alone, # p < 0.05 versus Normoxia Co-culture 1:1000; $ p < 0.05 versus respective condition islets alone.

For PDGF (Fig 5B), the TWA indicated significant effects of both the culturing conditions (p<0.0001) and the presence of ASC (p<0.0001), with no observed interaction. Comparing the secretomes derived from islets cultured alone to their baseline (normoxia-derived) secretome, a significant decrease in PDGF levels was evident in all secretomes (Šídák’s post-test, p<0.0001 for cytokines and high glucose, p=0.0003 for hypoxia + high glucose) except for the hypoxia-derived secretome. Similarly, significant decreased PDGF levels were observed in all secretomes (Šídák’s post-test, p<0.0001 for cytokines, high glucose, and hypoxia + high glucose) except for the hypoxia-derived one when comparing the secretomes from islets co-cultured with ASC to their baseline (normoxia-derived secretome). Further analysis of secretomes from islets co-cultured with ASC, compared to those from islets cultured alone, revealed an overall significant increase in PDGF levels in the co-culture secretomes (Šídák’s post-test, p=0.0006 for normoxia, p=0.0198 for cytokines, p=0.0097 for high glucose, p<0.0001 for hypoxia, p=0.0028 for hypoxia + high glucose).

For bFGF (Fig. 5C), the TWA demonstrated significant effects of the culturing conditions (p<0.0001), the presence of ASC (p<0.0001), and their interaction (p<0.0001). Comparing the secretomes derived from islets cultured alone to their baseline (normoxia-derived) secretome, a significant decrease in bFGF levels was only detected in the hypoxia-derived secretome (Šídák’s post-test, p=0.0393). When comparing the secretomes from islets co-cultured with ASC to their baseline (normoxia-derived secretome), significantly decreased bFGF levels were observed only in the high glucose-derived secretome (Šídák’s post-test, p<0.0001). In the comparison of various secretomes from islets cultured with ASC to those from islets cultured alone, an overall significant increase in bFGF levels was observed (Šídák’s post-test, p<0.0001 for normoxia, cytokines, hypoxia, and hypoxia + high glucose, p=0.0084 for high glucose).

For collagen I alpha 1 (Fig. 5D), the TWA demonstrated significant effects of the culturing conditions (p<0.0001), the presence of ASC (p<0.0001), and their interaction (p=0.0385). Comparing the secretomes derived from islets cultured alone to their baseline (normoxia-derived) secretome, a significant decrease in collagen I alpha 1 was observed in the cytokines (Šídák’s post-test, p=0.0023) and hypoxia + high glucose-derived secretomes (Šídák’s post-test, p=0.0400). When comparing the secretomes from islets co-cultured with ASC to their baseline (normoxia-derived secretome), significantly decreased collagen I alpha 1 levels were observed in the cytokines (Šídák’s post-test, p<0.0001) and high glucose-derived secretome (Šídák’s post-test, p=0.0023). When comparing various secretomes from islets cultured with ASC to those from islets cultured alone, an overall robust and significant increase in collagen I alpha 1 was observed (Šídák’s post-test, p<0.0001 for all).

For IL-1α (Fig. 5E), the TWA demonstrated significant effects of both culturing conditions (p=0.0004) and the presence of ASC (p=0.0002). When comparing the secretomes derived from islets cultured alone to their baseline (normoxia-derived) secretome, no changes in IL-1α were identified. Similarly, no changes were found when comparing the secretomes from islets co-cultured with ASC to their baseline (normoxia-derived secretome). In the comparison of various secretomes from islets cultured with ASC to those from islets cultured alone, despite a tendency towards an overall decrease, IL-1α levels were only significantly reduced in the cytokines-derived co-culture secretome (Šídák’s post-test, p<0.0174).

For IL-10 (Fig. 5F), the TWA showed significant effects of the culturing conditions (p<0.0001), the presence of ASC (p=0.0018), and their interaction (p=0.0028). Comparing the secretomes derived from islets cultured alone to their baseline (normoxia-derived) secretome, a significant decrease in IL-10 levels was only detected in the cytokines-derived secretome (Šídák’s post-test, p=0.0022). When comparing the secretomes from islets co-cultured with ASC to their baseline (normoxia-derived secretome), significantly decreased IL-10 levels were observed only in the high glucose-derived secretome (Šídák’s post-test, p=0.0277). In the comparison of various secretomes from islets cultured with ASC to those from islets cultured alone, despite a tendency towards an overall increase in IL-10 levels, a significant increase was only observed in the hypoxia + high glucose-derived secretome (Šídák’s post-test, p=0.0018).

#### 3.3.2. The concentration of key proteins in secretomes from rat islets with or without ASC

Figure 6 shows the analysis of key proteins involved in the pathways of interest within the secretome of rat islets cultured alone or with ASC. For VEGF (Fig. 6A), culturing conditions (p<0.0001), ASC presence (p<0.0001), and their interaction (p=0.0079) significantly influenced the TWA. When comparing islets cultured alone to their baseline (normoxia-derived) secretome, a general decrease in VEGF levels was observed (Šídák’s post-test, p<0.0001 for cytokines, p=0.0004 for high glucose, p=0.0056 for hypoxia, p=0.0056 for hypoxia + high glucose). Islets co-cultured with ASC exhibited a significant decrease in VEGF in all secretomes compared to their baseline (normoxia-derived secretome) (Šídák’s post-test, p<0.0001 for cytokines and high glucose, p=0.0002 for hypoxia + high glucose), except for the hypoxia-derived secretome. Further analysis of secretomes from islets cultured with ASC, compared to those from islets cultured alone, revealed a significant VEGF increase only in the hypoxia-derived secretome (Šídák’s post-test, p<0.0001).

**Fig. 6.**
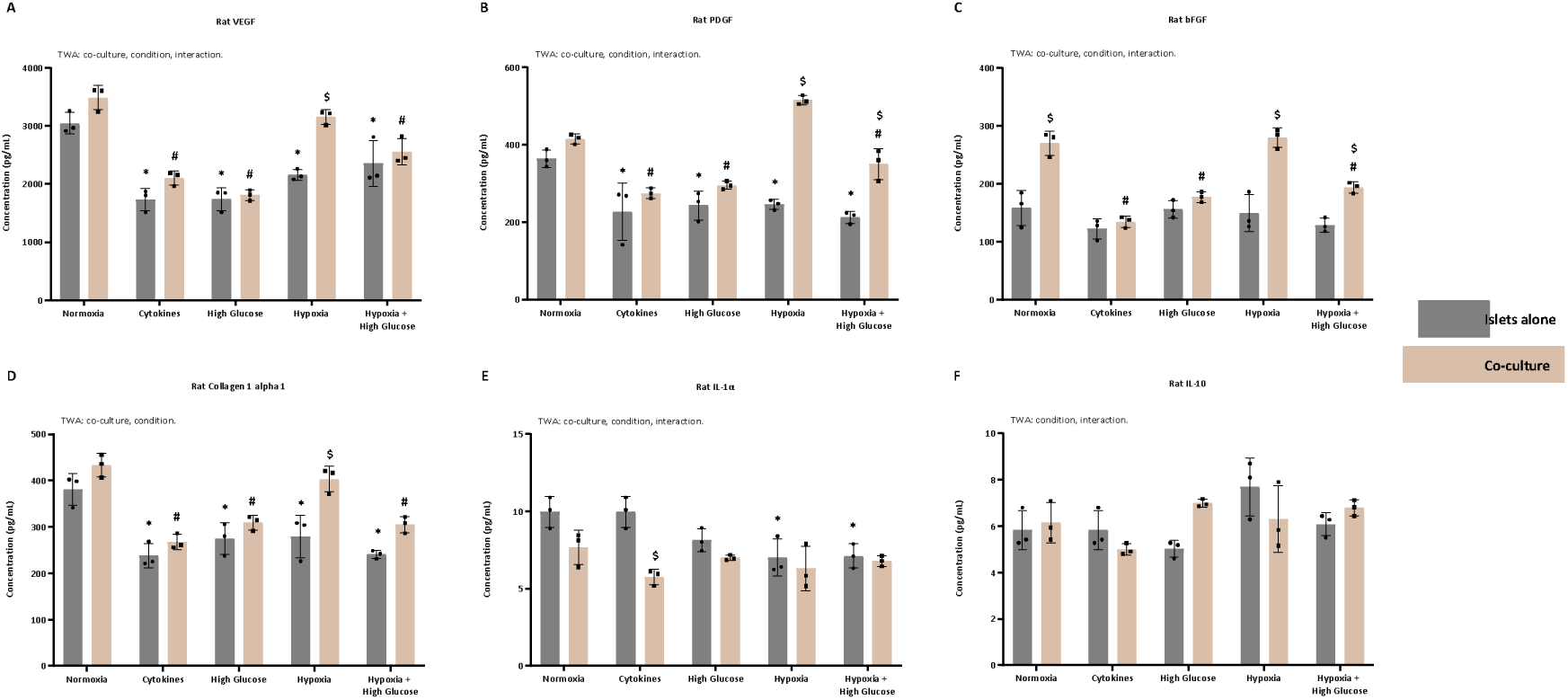
Modulation of key paracrine factors involved in pathways of interest on the rat secretome upon co-culturing. Secretion of (A) vascular endothelial growth factor (VEGF), (B) platelet-derived growth factor (PDGF), (C) basic fibroblast growth factor (b-FGF), (D) collagen I alpha I, (E) interleukin 1 alpha (IL-1α), and (F) IL-10 after 72 hours co-culturing in standard conditions (21% O2). Data represent mean values ±0standard deviation of 3 pooled samples measured in triplicate. Two-way ANOVA (TWA) followed by Šídák’s multiple comparison test, * p < 0.05 versus Normoxia islets alone, # p < 0.05 versus Normoxia Co-culture 1:1000; $ p < 0.05 versus respective condition islets alone.

For PDGF (Fig. 6B), the TWA demonstrated significant effects of the different culturing conditions (p<0.0001), the presence of ASC (p<0.0001), and their interaction (p<0.0001). Comparing the secretomes from islets cultured alone to their baseline (normoxia-derived) secretome, an overall significant decrease in PDGF levels was noted (Šídák’s post-test, p=0.004 for cytokines, p=0.0018 for high glucose, p=0.0022 for hypoxia, p=0.0001 for hypoxia + high glucose). When comparing secretomes from islets co-cultured with ASC to their baseline (normoxia-derived secretome), again all secretomes exhibited significantly reduced PDGF levels (Šídák’s post-test, p=0.000 for cytokines, p=0.0019 for high glucose, p=0.0117 for hypoxia), except for the hypoxia + high glucose-derived secretome. Further analysis of secretomes from islets cultured with ASC, compared to those from islets cultured alone, revealed a significant increase in PDGF levels in the co-culture secretomes derived from hypoxia (Šídák’s post-test, p<0.0001) and hypoxia + high glucose culturing (Šídák’s post-test, p=0.0004).

For bFGF (Fig. 6C), the TWA demonstrated significant effects of the culturing conditions (p<0.0001), the presence of ASC (p<0.0001), and their interaction (p<0.0001). Comparing the secretomes derived from islets cultured alone to their baseline (normoxia-derived) secretome, no changes in bFGF levels were identified. When comparing the secretomes from islets co-cultured with ASC to their baseline (normoxia-derived secretome), significantly decreased bFGF levels were observed in the cytokines (Šídák’s post-test, p<0.0001), high glucose (Šídák’s post-test, p=0.0001), and hypoxia + high glucose-derived secretome (Šídák’s post-test, p=0.0011). In the comparison of various secretomes from islets cultured with ASC to those from islets cultured alone, significantly increased bFGF levels were observed in the normoxia, hypoxia (Šídák’s post-test, p<0.0001), and hypoxia + high glucose-derived secretome (Šídák’s post-test, p=0.0063).

For collagen I alpha 1 (Fig. 6D), the TWA revealed significant effects of both culturing conditions (p<0.0001) and the presence of ASC (p<0.0001). Comparing the secretomes derived from islets cultured alone to their baseline (normoxia-derived) secretome, a significant decrease in collagen I alpha 1 was observed in all secretomes (Šídák’s post-test, p<0.0001 for cytokines and hypoxia + high glucose, p=0.0015 for high glucose, p=0.0026 for hypoxia). When comparing secretomes from islets co-cultured with ASC to their baseline (normoxia-derived secretome), significantly decreased collagen I alpha 1 levels were observed in all secretomes (Šídák’s post-test, p<0.0001 for cytokines, p=0.0003 for high glucose, p=0.0002 for hypoxia + high glucose), except for the hypoxia-derived one. In the comparison of various secretomes from islets cultured with ASC to those from islets cultured alone, significantly increased collagen I alpha 1 was observed only in the hypoxia-derived secretome (Šídák’s post-test, p=0.0003).

For IL-1α (Fig. 6E), the TWA indicated significant effects of the culturing conditions (p<0.0001), the presence of ASC (p=0.0054), and their interaction (p=0.0097). Comparing the secretomes derived from islets cultured alone to their baseline (normoxia-derived) secretome, a significant decrease in IL-1α was observed in the hypoxia (Šídák’s post-test, p=0.0101) and hypoxia + high glucose-derived secretomes (Šídák’s post-test, p=0.0134). In the comparison of secretomes from islets co-cultured with ASC to their baseline (normoxia-derived secretome), no significant changes in IL-1α levels were identified. When comparing various secretomes from islets cultured with ASC to those from islets cultured alone, despite a tendency towards an overall decrease, IL-1α levels were only significantly reduced in the cytokines-derived co-culture secretome (Šídák’s post-test, p=0.0002).

For IL-10 (Fig. 6F), the TWA revealed significant effects of the culturing conditions (p=0.0315) and interaction (p=0.0130). When comparing the secretomes derived from islets cultured alone to their baseline (normoxia-derived) secretome, no significant changes in IL-10 levels were detected. Similarly, no significant changes were identified when comparing the secretomes from islets co-cultured with ASC to their baseline (normoxia-derived secretome). Additionally, when comparing various secretomes from islets cultured with ASC to those from islets cultured alone, no modulation of IL-10 was found.

### 3.4. In vitro functional properties of islets (with or without ASC) secretome

Most analyzed secretomes contained proteins associated with angiogenesis, ECM composition, and immune responses. Next, we aimed to investigate *in vitro* the effects of the secretomes on angiogenesis, ECM composition, and immune responses using the TFA, the Picrosirius Red staining, and the MLR followed by an antibody-mediated CDC.

#### 3.4.1. The proangiogenic potential of human secretome

Fig. 7A illustrates the analysis of branching points resulting from the TFA with HUVEC incubated with the various secretomes from human islets alone or in co-culture with ASC under various culturing conditions. The TWA revealed significant effects of both culturing conditions (p<0.0001) and the presence of ASC (p<0.0001), with no interaction observed. Comparing secretomes derived from islets alone under different culturing conditions to their baseline (normoxia-derived) secretome, we observed a significant increase in branching points when HUVEC were exposed to secretomes from hypoxia (Šídák’s post-test: p=0.0005) and hypoxia + high glucose (Šídák’s post-test, p<0.0001). For co-cultured islets, the secretome from exposure to cytokines significantly decreased the number of branching points (Šídák’s post-test, p=0.0033), while the secretomes of hypoxia (Šídák’s post-test, p=0.0126), and hypoxia + high glucose (Šídák’s post-test, p=0.0060) significantly increased branching points formed by HUVEC as compared to the normoxia secretome of co-cultured islets. Notably, comparing the secretomes from all conditions from islets cultured alone with islets cultured with ASC, only the normoxia-derived co-culture secretome significantly enhanced the number of branching points formed by HUVEC as compared to the branching points after incubation with the islet alone secretome (Šídák’s post-test: p=0.0039).

**Fig. 7.**
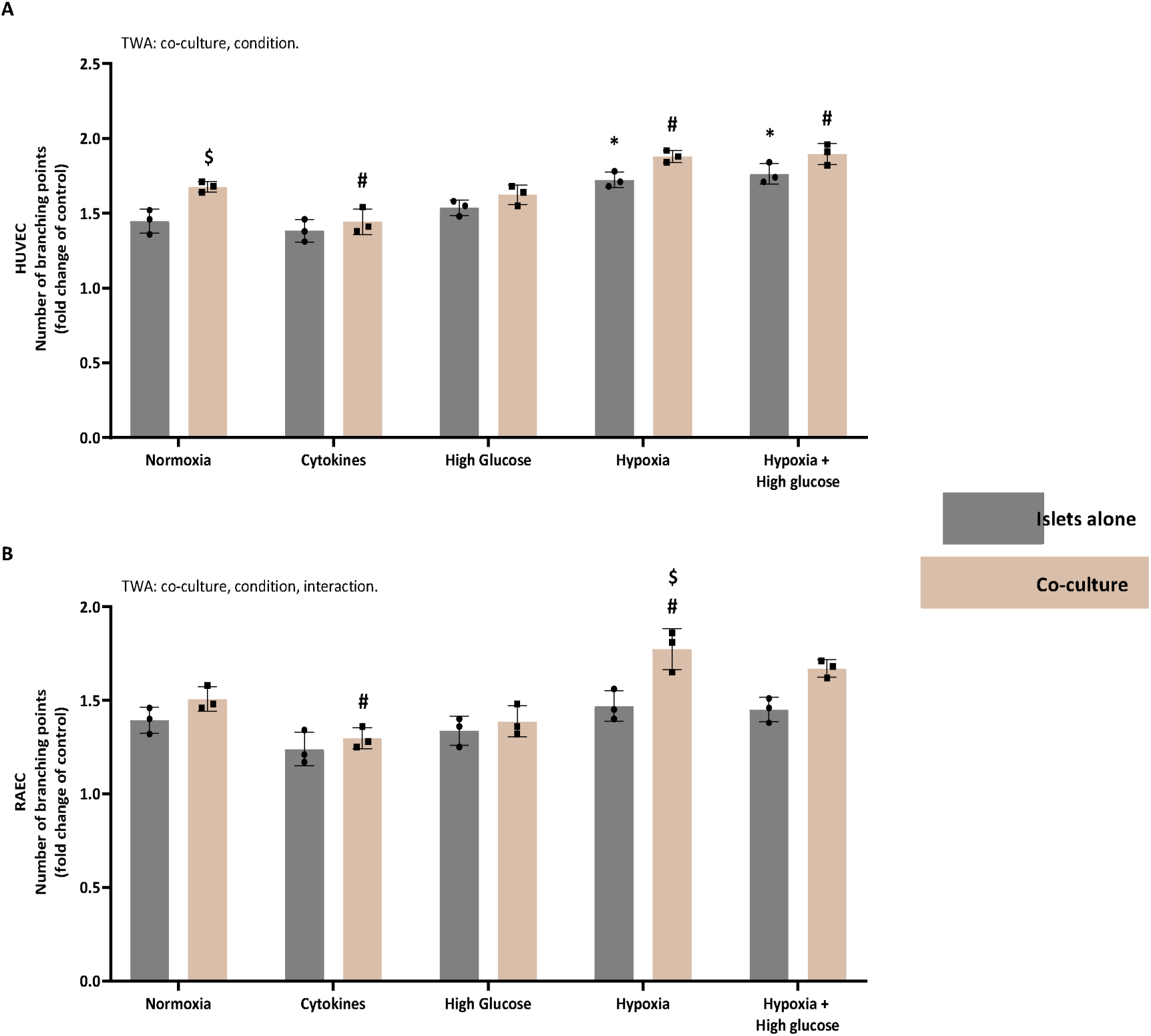
Co-culturing with ASC enhance the proangiogenic potential of the secretome. Quantification of the branching points formed by (A) human and (B) rat tube formation assay. HUVEC (Human Umbilical Vein Endothelial Cells) or RAEC (Rat Aortic Endothelial Cells) were exposed to the various secretomes derived from human or rat islets or co-culture 1:1000 for 20 hours (n = 3). The number of branching points was normalized to the control (CMRL (−); set at 1. Data are represented as individual values and mean ± standard deviation. Statistical significance was assessed using two-way ANOVA (TWA) and Šídák’s multiple comparison test, * p < 0.05 versus Normoxia islets alone, # p < 0.05 versus Normoxia Co-culture 1:1000, $ p < 0.05 versus respective condition islets alone.

#### 3.4.2. The proangiogenic potential of rat secretome

Fig. 7B depicts the analysis of branching points resulting from the TFA with RAEC, utilizing secretomes from rat islets alone or in co-culture with rat ASC under various culturing conditions. The TWA demonstrated significant effects of both the culturing conditions (p<0.0001) and the presence of ASC (p<0.0001). Importantly, an interaction between culturing conditions and the presence of ASC was observed (p=0.038), indicating a distinct impact of ASC under different culturing conditions. When comparing different culturing conditions for islets alone, none of the secretomes affected the number of branching points differently than their baseline (normoxia-derived) secretome. In contrast, for islets cultured with ASC, secretomes derived from cytokines exposure (Šídák’s post-test, p=0.039) significantly decreased the number of branching points compared with the normoxic secretome, while hypoxia (Šídák’s post-test, p=0.0048) significantly increased the number of branching points compared to the normoxic secretome. Post-testing also showed that only using hypoxia (Šídák’s post-test, p=0.0012) and high glucose + hypoxia (Šídák’s post-test, p=0.0271) secretomes there was an increase in the number of branching points with the secretome from islets with ASCs compared to the secretome of only islets.

#### 3.4.3. The potential of human secretome to modulate collagen deposition

Figure 8A presents the analysis of collagen deposition by human fibroblasts following incubation with secretomes from human islets either alone or in co-culture with human ASC under diverse culturing conditions. The TWA indicated significant effects of both culturing conditions (p<0.0001) and the presence of ASC (p<0.0001), with no observed interaction between culturing conditions and ASC presence. For islets alone, compared to normoxic culturing, the cytokines secretomes (Šídák’s post-test, p<0.0001), high glucose secretomes (Šídák’s post-test, p<0.0001), and high glucose + hypoxia secretomes (Šídák’s post-test, p<0.0296) induced decreased collagen deposition, while the hypoxia secretome (Šídák’s post-test, p<0.0327) increased collagen deposition compared to the normoxic secretome. Similarly, for islets with ASC, compared to normoxic culturing, both cytokine secretomes (Šídák’s post-test, p<0.0001) and high glucose secretomes (Šídák’s post-test, p<0.0001) induced decreased collagen deposition, while hypoxia (Šídák’s post-test, p<0.0071) increased collagen deposition compared to the normoxic secretome. Post-testing also revealed that under all conditions, the secretome of islets co-cultured with ASC induced significantly more collagen deposition compared to islets alone (Šídák’s post-test, p=0.0327 for cytokines; p=0.0019 for high glucose; p=0.0216 for hypoxia; p=0.0013 for hypoxia + high glucose).

**Fig. 8.**
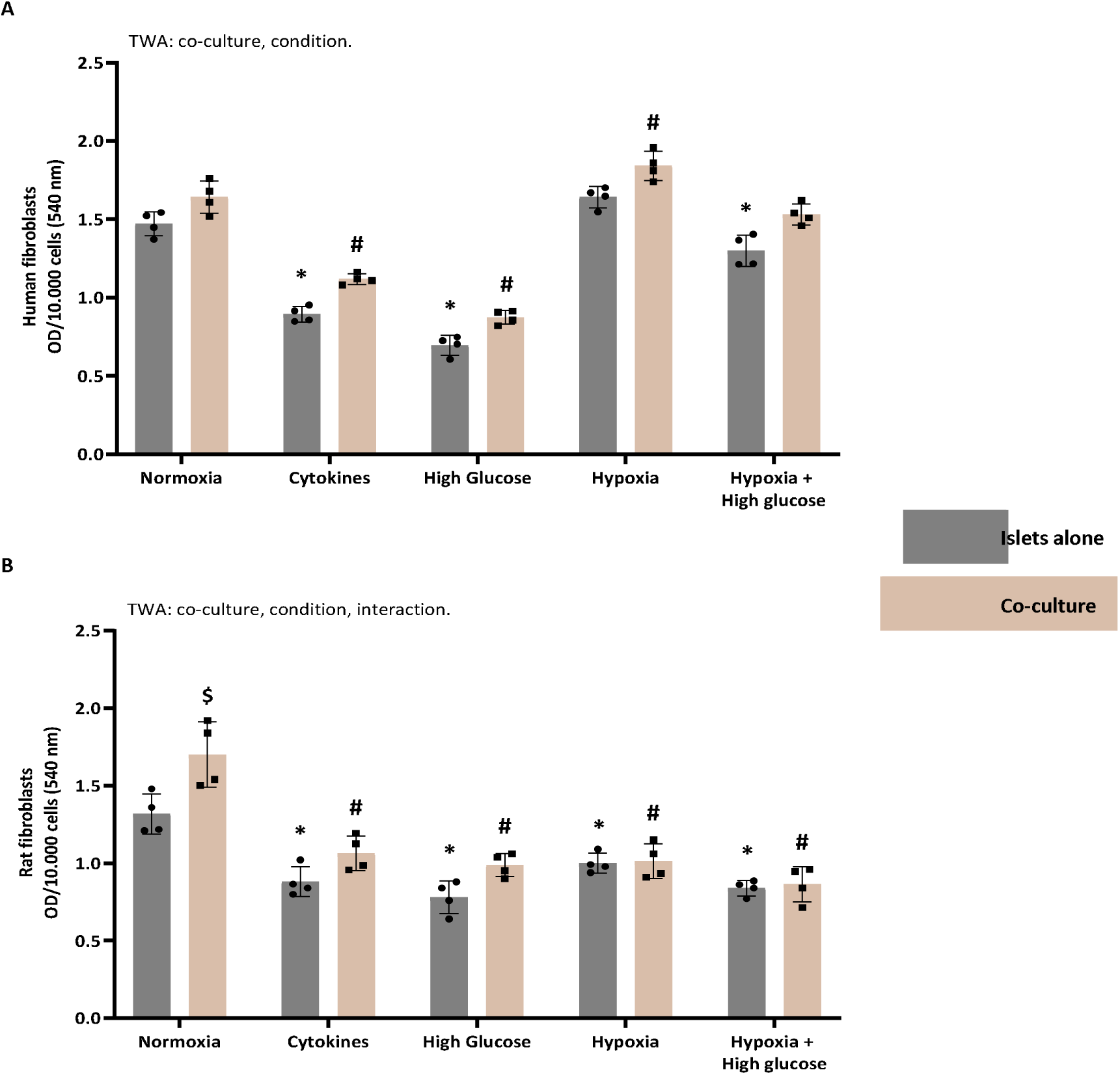
Enhanced collagen deposition by fibroblast exposed to co-culture secretome. Spectrophotometric analysis of the Sirius-Red-stained. (A) Human and (B) rat fibroblasts exposed to various secretomes derived from human or rat islets or co-culture 1:1000 for 72 hours (n = 4). The data was normalized to the control (STD-M (−); set at 1. Data are represented as individual values and mean ± standard deviation. Statistical significance was assessed using two-way ANOVA (TWA) and Šídák’s multiple comparison test, * p < 0.05 versus Normoxia islets alone, # p < 0.05 versus Normoxia Co-culture 1:1000. $ p < 0.05 versus respective condition islets alone.

#### 3.4.4. The potential of rat secretome to modulate collagen deposition

Fig. 8B displays the analysis of collagen deposition by rat fibroblasts following incubation with secretomes from rat islets either alone or in co-culture with rat ASC under diverse culturing conditions. The TWA indicated significant effects of both culturing conditions (p<0.0001) and the presence of ASC (p=0.0001), with an observed interaction between culturing conditions and ASC presence (TWA, p=0.0188). For islets alone, all secretomes significantly decreased collagen deposition compared to normoxic culturing (Šídák’s post-test, p=0.0001 for cytokines; p<0.0001 for high glucose; p=0.0065 for hypoxia; p<0.0001 for hypoxia + high glucose). Similarly, significant decreased collagen deposition was observed for all secretomes of islets with ASC compared with normoxia (Šídák’s post-test, p<0.0001 for all conditions). Post-testing also revealed that only under normoxia the secretome of islets co-cultured with ASC induced significantly more collagen deposition than islets alone (Šídák’s post-test, p<0.0007).

#### 3.4.5. The immunomodulatory potential of human secretome

Fig. 9A illustrates the survival percentage of human PBMC after the antibody-mediated CDC assay. The effectiveness of the antibody-mediated CDC assay is evident from a significant decline (p < 0.0001) in PBMC survival after incubation with MLR supernatant (only alloantibodies without secretome) compared to resting PBMC exposed solely to STD-M (+) (100 % survival, data not shown in the graph).

**Fig. 9.**
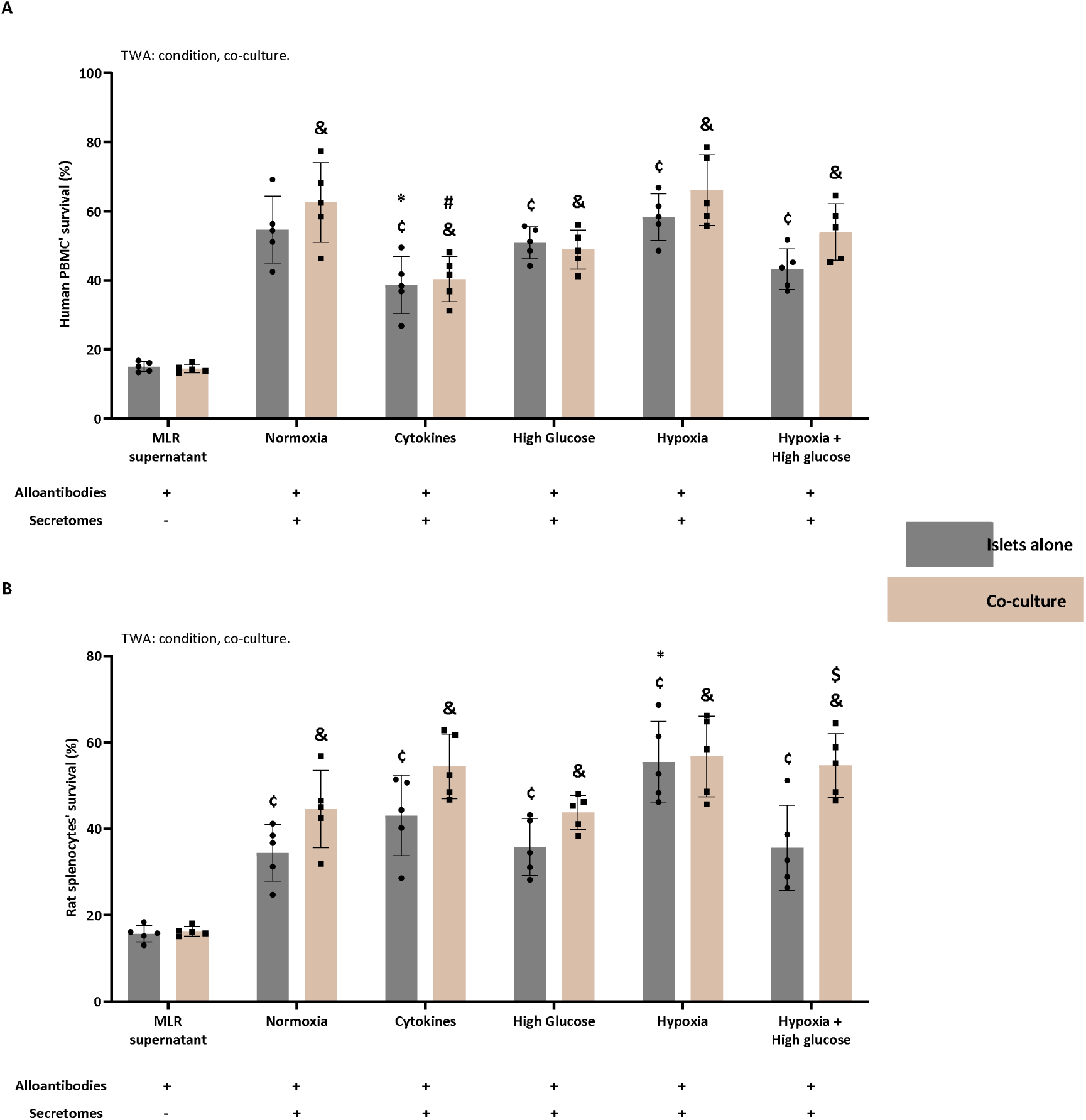
Enhanced capacity of the co-culture secretome to modulate antibody-mediated immune responses. The effect of various secretomes derived from (A) human and (B) rat islets and co-cultures 1:1000 on humoral alloimmunity, evaluated using a mixed lymphocyte reaction (MLR) followed by an antibody-mediated complement-dependent cytotoxicity (CDC) assay. The outcome is expressed as the percentage of human Peripheral Blood Mononuclear cells (PBMC) or rat splenocytes that survived the exposure to the alloantibodies (n = 5 for human and rat). CMRL (−) was used as a control and together with the MLR supernatant it shows the efficacy of the antibody-mediated CDC assay. Cell survival was normalized to the control (CMRL (−); set to 100). Data are represented as mean ± standard deviation. Statistical significance was assessed using two-way ANOVA (TWA) and Šídák’s post hoc test. ¢ versus MLR supernatant islets alone, & versus MLR supernatant co-culture, * p < 0.05 versus Normoxia islets alone, # p < 0.05 versus Normoxia Co-culture 1:1000, $ p < 0.05 versus respective condition islets alone.

The TWA indicated significant effects of both culturing conditions (p<0.0001) and the presence of ASC (p=0.0287). MLR supernatants, generated in the presence of secretomes derived from both islets alone or co-cultured islets under various conditions, induced increased PBMC survival (Šídák’s post-test, p<0.0001) compared to MLR supernatant only (containing no secretome). The PBMC survival after incubation with the alloantibodies generated in the MLR in the presence of the different secretomes of the islets alone was compared to the PBMC survival after incubation with alloantibodies generated in the MLR in the presence of the normoxic secretome of the islets alone. This showed that the PBMC survival of the human cytokines-derived islet secretome significantly decreased compared to PBMC survival of the normoxia-derived islet secretome (Šídák’s post-test, p=0.0261). When comparing the different secretome conditions to the normoxic secretome from the co-cultured islets, the PBMC survival after incubation of alloantibodies generated in the presence of the cytokines-derived co-culture secretome was significantly decreased (Šídák’s post-test, p=0.0004). No significant effect was found when comparing the islets alone with the co-culture per condition.

#### 3.4.6. The immunomodulatory potential of rat secretome

Figure 9B shows the survival percentage of rat splenocytes after the antibody-mediated CDC assay. The efficiency of the assay was shown by the significant decline (p < 0.0001) in splenocyte survival after incubation with MLR supernatant (containing only alloantibodies and no secretome) when compared with splenocytes exposed solely to STD-M (+) (100 %; data not shown in the graph).

The TWA indicated that there was an effect of the culturing conditions (TWA: p<0.0001) and the presence of ASC (TWA: p<0.0001). MLR supernatants generated in the presence of all secretomes from both islets or co-cultured islets significantly increased splenocyte survival compared to the MLR supernatant only (containing only alloantibodies). When comparing the different secretome conditions to the normoxic secretome of the islets alone, the splenocyte survival after incubation with the hypoxia-derived islets secretome was significantly increased (Šídák’s post-test, p=0.0010). When comparing the different secretome conditions to the normoxic secretome of the co-cultured islets, there was no significant difference. Post-testing also showed a significant increase in the survival of splenocytes exposed to the MLR supernatant generated in the presence of hypoxia + high glucose-derived secretomes from islets with ASC (Šídák’s post-test, p=0.0038) compared to the ones exposed to its counterpart from islets cultured alone.

## 4. Discussion

Despite the positive functional effects of ASC co-culture or co-transplantation with islets observed in previous studies [6, 12, 13, 16, 17], the specific mechanisms of how ASC co-culture/co-transplantation influences islets are still poorly understood. This study delves into the secretome during co-culturing of islets and ASC, aiming to understand the factors secreted and their potential effects on islet function and survival. We hypothesized that co-culturing islets with ASC produces a secretome containing factors capable of enhancing the islets’ microenvironment. This enhancement involves stimulating angiogenesis, promoting the deposition of ECM components, and providing immune modulation. Our data show that co-culturing islets with ASC under different conditions, such as normoxia, hypoxia, or high glucose, influenced the secretome. Many pathways associated with angiogenesis, ECM, and immune responses were affected, which may explain the beneficial effects of the ASC co-culture/co-transplantation on islet function and survival following PIT. This research contributes to a deeper understanding of the impact of ASC co-culture on pancreatic islets.

Islets co-cultured with ASC manifest a distinctive secretion fingerprint compared to islets cultured alone. This was evident from the baseline, normoxia-derived secretomes, and functional analyses of both human and rat models. Cultured alone under normoxia, islets secreted proteins that participate in pathways of protein homeostasis and basal cell responses (e.g., response to xenobiotic stimulus, regulation of proteolysis, protein maturation, and others). However, the introduction of ASC induces specific enhancements and alterations. In the normoxia-derived secretome of co-cultured human islets, there is a notable increase in pathways linked to energy generation, protein homeostasis, and ECM organization. PDGF, bFGF, and Collagen I alpha 1 secretion were also increased after the co-culture of islets with ASC compared to the culture of islets alone. These latter findings correlated with the functional assays in which co-culture with ASC increased the number of branching points in HUVEC as well as the deposition of collagen by fibroblasts. Collectively, this indicates a secretome that can influence the microenvironment to support vital processes such as energy production, protein balance, structural organization, and angiogenesis. These processes are crucial not only for the health and optimal functioning of islets but also hold the potential for enhancing adaptability in transplantation scenarios. The normoxia-derived rat islet secretome showed a similar profile to human islets, especially in pathways linked to protein homeostasis (e.g., protein maturation, regulation of proteolysis). Once co-cultured with ASC, rat islets under normoxic circumstances showed enhancement in the pathway profile related to protein homeostasis, and the presence of pathways of responses to xenobiotic stimuli, wounding, and metal ions, but also pathways related to the ECM were enriched. This is supported by the functional assay, showing increased collagen deposition by fibroblasts following incubation with the secretome from this condition. Supported by the functional assays (MLR and TFA), pathways associated with the immune systems and angiogenesis were not enriched in this secretome.

Differences were also observed when comparing the secretomes from islets co-cultured with or without ASC in response to cytokines. When cultured alone, human and rat islets shared metabolic and regenerative responses to cytokines stimulation, with enrichments in energy generation, protein maturation, and response to wounding, which may indicate increased metabolic activity, enhanced synthesis of functional proteins, and an activated regenerative response [30–32]. However, species-specific differences arose in carbohydrate metabolism, lipid-related processes, and specific cellular responses, reflecting multifaceted influences from evolutionary and physiological factors [33–35]. Conversely, co-cultured islets with ASC displayed unique enrichments, particularly in the cytokine-derived secretome from human islets, showing enrichment in pathways related to protein maturation, peptide metabolic processes, and ECM organization. Functional assays revealed a significant increased potential for collagen deposition, indicating that the pathways associated with ECM organization may be increased. This may create a more supportive microenvironment for islet health, which is crucial post-isolation and transplantation when islets face challenges with a damaged ECM [36–38]. Co-cultured rat islets under cytokine exposure demonstrated distinct responses, including responses to xenobiotic stimuli and enrichment in protein folding pathways, indicating increased adaptability to foreign substances and optimized mechanisms for proper protein maturation. These adaptations may contribute to improved graft viability, enabling transplanted islets to withstand better the stresses associated with the transplantation process, similar to the benefits observed in human islets.

In the context of high glucose-derived secretomes, for islets cultured alone (human and rat), the enrichment of pathways related to protein maturation and folding may result from an adaptive response to the elevated glucose levels [39–41]. This includes an increased demand for proper protein folding and cellular damage repair [40, 42]. Notably, when co-cultured with ASC, human islets exposed to high glucose enriched their secretome with proteins associated with ECM pathways, such as collagen I alpha 1. This is also apparent from our functional assay, which showed increased collagen deposition under the influence of the high glucose co-culture secretome compared to the secretome derived from islet culture alone. Concurrently, co-cultured rat islets also experience further enrichment in protein maturation and folding pathways. These adaptations collectively contribute to a more supportive microenvironment, suggesting that co-culturing with ASC may positively impact islet functionality in high glucose conditions.

Under hypoxia, human and rat islets cultured alone displayed a secretome enriched in pathways that highlight metabolism homeostasis and energy generation (with enriched pathways such as generation of precursor metabolites and energy, cellular respiration, peptide metabolic process, and others), showing that adaptive responses to low oxygen levels are significantly reflected in the islets’ secretion signature. Specifically, the enrichment in pathways related to metabolic homeostasis indicates an effort by the islets to maintain a balance in cellular processes, possibly to sustain essential functions despite reduced oxygen availability. Additionally, the emphasis on energy generation pathways suggests an increased demand for energy production, potentially to support cellular activities and cope with the challenges imposed by hypoxic conditions. When co-cultured with ASC, the pathways enrichment of islets exposed to hypoxia (both human and rat) shows enhancement in pathways such as response to reactive oxygen species, ECM, and structure organization. The functional assays showed that hypoxia-derived secretomes from rat co-cultures exhibited a higher proangiogenic potential in comparison to those from islets cultured alone in the same condition. Similarly, human co-cultures demonstrated increased collagen secretion potential when compared to the secretome of islets cultured alone under hypoxia. These findings suggest that for these specific hypoxia-derived secretomes, ASC not only contribute to creating a microenvironment that enhances adaptability and resilience to low oxygen levels but also stimulates functional improvements. These improvements could offer advantages in terms of promoting angiogenesis and facilitating tissue repair, which are crucial for the overall health and performance of transplanted islets.

Finally, the exposure of human and rat islets to the combination of hypoxia and high glucose conditions showed that in such extreme stress conditions, islets enrich their secretome with factors linked mainly to pathways of protein and cellular homeostasis. When co-cultured with ASC, the secretome of islets (human and rat) exposed to hypoxia and high glucose displayed an overall enrichment in pathways of cellular response, protein homeostasis, and ECM organization. Functionally, human co-cultured secretomes displayed increased capacity to stimulate collagen deposition. Rat co-cultured secretome, in turn, showed enhanced proangiogenic and also immunomodulatory potential. In the context of this dual stress, co-culturing may be instrumental in enhancing the resilience and functionality of both human and rat islets, mitigating the detrimental effects of extreme stress.

Our pathway enrichment analysis noted that the top 5 most enriched pathways in the secretome of islets cultured alone consistently appeared within the top 100 pathways in the secretome of islets cultured with ASC. This uniformity implies a robust and stable composition of the islet secretome, indicating the fundamental role of certain pathways in islet function under specific conditions. Thus, the presence of ASC does not disrupt the core enriched pathways; instead, it introduces additional pathways. This suggests a potential synergistic or modulatory effect of ASC on the islet secretome, indicating that the co-culture may enhance the complexity or functionality of the islet secretome by activating or influencing supplementary pathways. An alternate explanation is that co-culturing islets with ASC may not significantly alter the islets’ secretome itself. Instead, the observed differences in the secretome between islets co-cultured with ASC and islets cultured alone may arise from the influence of the ASC secretome. Further research is warranted to substantiate and identify the source of these additional pathways induced by ASC co-culturing. Such investigations may yield more insights into the broader biological interactions and functions associated with islets and ASC, potentially paving the way for new research and therapeutic development.

In conclusion, studying the secretome is a valuable tool to gain profound insights into the benefits of islets co-cultured with ASC. The observed enrichments in crucial pathways such as cellular response, protein homeostasis, and ECM organization, coupled with functional enhancements, offer a comprehensive understanding of how co-culturing with ASC enhances islet function and survival. This unique perspective not only unveils the adaptive responses of islets to stressors but also underscores the transformative impact of ASC co-culture. By unravelling the molecular signatures of secretomes, this study contributes to scientific understanding and lays the foundation for developing innovative strategies to optimize islet performance, particularly after transplantation.

## Supporting information

Supplementary tables Human

Supplementary tables Rat

## Acknowledgments

We thank dr. Marijke M. Faas for all the help and inspiring discussions regarding this work. This research was supported by the Dutch Diabetes Research Foundation [2019.81.001].

## Author contributions

All authors were actively engaged in the study. E.P.M contributed to the conception, design, data collection and assembly, data analysis and interpretation, manuscript writing, and final approval. B.J.H. was involved in the design and final approval of the manuscript. M.A.E. provided study material and approved the final manuscript. A.M.S. contributed to the conception, design, and final approval of the manuscript, as well as providing financial and administrative support.

