## Supplementary figure 1 for "Secretome analysis of human and rat pancreatic islets co-cultured with adipose-derived stromal cells reveals a signature with enhanced regenerative capacities"

**Supplementary Figures**

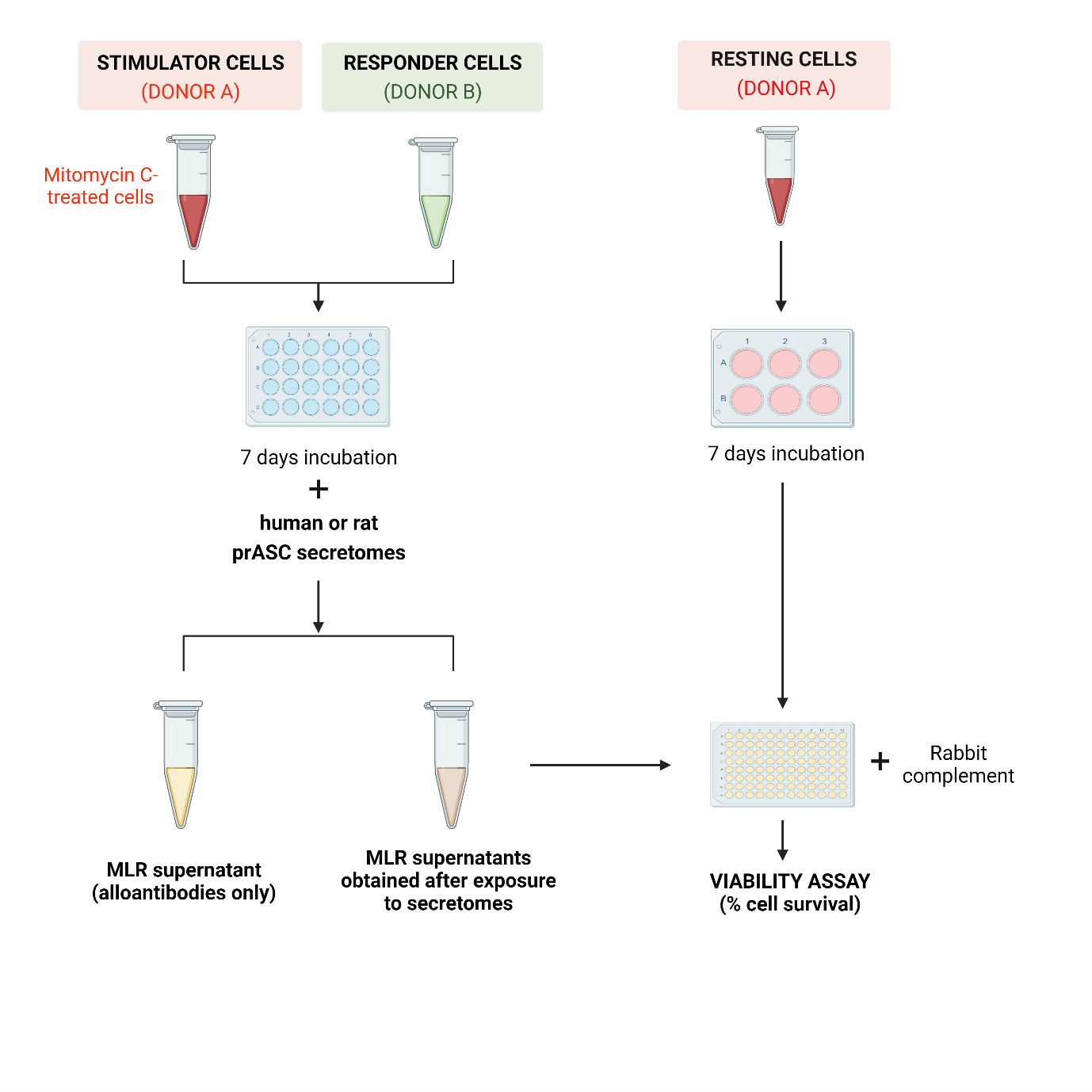

**Supplementary Fig 1.** Illustrative representation of the two-way mixed lymphocyte reaction (MLR) followed by an antibody-mediated cell dependent cytotoxicity assay (CDC) protocol used to evaluate the various human and rat prASC secretomes' capacity to modulate antibody-mediate immune responses. Created with BioRender.com.
